# A high-resolution aerial camera survey of Uganda’s Queen Elizabeth Protected Area improves detection of wildlife and delivers a surprisingly high estimate of the elephant population

**DOI:** 10.1101/2023.02.06.525067

**Authors:** Richard H Lamprey, Michael Keigwin, Charles Tumwesigye

## Abstract

The Queen Elizabeth Protected Area (QEPA) hosts some 60% of Uganda’s elephants and large populations of buffalo, hippopotamus and Uganda kob. However, the area is subjected to continued poaching, livestock incursions, animal disease and invasive plant species. Population estimates derived from aerial observers have shown great variability, and therefore trends are hard to discern. The Uganda Wildlife Authority (UWA) reguires precise wildlife population estimates to guide conservation actions. The aim of this study is to provide accurate and precise baseline estimates for wildlife populations of QEPA using aerial imagery and high sampling intensity, and to improve survey methods for determining future trends. High-resolution cameras, orientated at 45°, captured images along sample strips to left and right of the aircraft. Transects at 1 km separation gave a high sampling intensity of 28% to improve precision. We captured 43000 images for visual enumeration of 13 wildlife species. Population estimates (Ŷ), standard errors (SE) and relative margins of error (RME =95% c.l/Ŷ) were determined using ‘Jolly II’, Marriott and bootstrapping methods. With improved detection using imagery the QEPA elephant population is estimated at 4 711 ± 1106 (95% c. I), which is 62% higher than the previous observer-based estimate of 2900 in 2014, and the highest since counts began in the 1960s. The survey achieved an RME for elephants of 23%, making this one of the most precise counts for any similar-sized elephant area in Africa. The buffalo, Uganda kob and hippopotamus populations have stabilized at ‘safe’ levels but remain well below the high point of the mid-1990s; the high density of bone patches indicate high mortality, attributable to disease and to the continued invasion of the unpalatable grass species <u>Imperata cylindrica</u>, which now covers 50% of the grassland area. Our methods indicate that methods and sample parameters prescribed by international elephant counting standards can be revised and improved using aerial cameras, especially for smaller survey areas.

**Short Summary Text:** Uganda’s Queen Elizabeth Protected Area (QEPA) continues its recovery since the decimation of wildlife by militias in the 1970s, but it is challenging to track wildlife trends due to the variability in population estimates. To improve accuracy and precision, we conducted the first aerial count of QEPA using high-resolution imaging. This approach has indicated that the elephant population is at its highest levels since surveys began in the 1960s. Meanwhile, buffalo, topi and Uganda kob are stagnating below previous 1990s levels. We suggest the use of camera systems in future surveys to improve the accuracy and precision of counts, in order to guide effective management.

## 1. INTRODUCTION

This paper reports on the methods and results of an aerial wildlife survey of Queen Elizabeth Protected Area (QEPA) in the Albertine Rift of Uganda, conducted in 2018. In this count, human rear-seat-observers (RSOs) were replaced with ‘oblique-camera-count’ (OCC) imaging systems (Lamprey, Ochanda, Brett, Tumwesigye, & Douglas-Hamilton, 2020). Previous aerial counts in QEPA using RSOs have not been sufficiently precise to determine how wildlife populations have been impacted by perturbations, especially since the turn of the millennium. These ecosystem threats include wildlife disease, livestock invasions, expansion of human enclaves, the spread of invasive plant species, and resurgent poaching, due in large part to the COVID-19 pandemic. QEPA’s new responsive and adaptive management team urgently requires more accurate and precise information on the status of wildlife populations and distributions.

QEPA is a conservation unit of 2842 km^2^ comprised of Queen Elizabeth National Park, established in 1953 and named in commemoration of the Queen of England the year before, and its smaller contiguous wildlife reserves of Kyambura and Kigezi. It protects lake and savannah habitats in one of the world’s most biodiverse areas, the Greater Virunga Landscape (GVL) of northern DR Congo and Western Uganda (Plumptre, Kujirakwinja, Treves, Owiunji, & Rainer, 2007). With its backdrop of the Rwenzori Mountains and its contiguous Lakes Edward and George, QEPA is renowned for its spectacular wetlands, grasslands and forests, which support high densities of elephants *(Loxodonta africana)*, buffalo *(Syncerus coffer)*, hippopotamus *(hippopotamus amphibius)*, and Uganda kob *(Kobus kob* ssp *thomasi)*. The area has probably the highest diversity of birds in any protected area in Africa, with over 600 bird species recorded (BirdLife_International, 2021).

In the 1960s, the terrestrial large mammal biomass of QEPA was considered the highest in Africa, and possibly the world (Coe, Cumming, & Phillipson, 1976; Field & Laws, 1970; Petrides & Swank, 1965). However, there are relatively few large mammal species, a circumstance attributed to habitat alteration and hunting by the many agricultural and fishing communities inhabiting the area in the 1800s and early 1900s (Spinage, 1970). Eland, impala, zebra, hartebeest and giraffe, present in other Uganda protected areas in western Uganda have not historically been reported in QEPA; it is assumed that if they had previously existed there, they were either hunted out or extirpated by the rinderpest pandemic of 1890. Between 1910 and 1930 severe outbreaks of sleeping sickness prompted the relocation of farming, fishing and pastoral communities away from lakeshores, which opened the area to immigration of herds of elephants and buffalo from other ‘Great Lakes’ areas. The complex history of QEPA, declared Uganda’s first national park in 1952, is reflected in its convoluted boundary that takes in human fishing village ‘enclaves’ and public road systems; people and wildlife are in close contact across all habitats (Lamprey et al., 2000).

Monitoring of wildlife in QEPA is further complicated by the park’s contiguous and permeable boundary with the Parc National des Virungas (PNV) in neighbouring Democratic Republic of Congo, which adjoins QEPA along a 20 km stretch of the Ishasha River south of Lake Edward (the ‘Ishasha Sector^7^), and through a thin strip of woodland to the north of the lake. The large mammals of Ishasha, especially elephant, buffalo and topi *(Damaliscus lunatus* ssp *jimela)*, may move unimpeded between the ‘Lake Edward flats’ of Q.ENP and Virunga (Mubalama, 2000). With different wildlife management agencies across this Anglophone/ Francophone border, only recently has it been possible to coordinate aerial surveys in QEPA and Virunga to understand wildlife distributions across the entire QEPA/PNV ecosystem (Plumptre, Kujirakwinja, Moyer, Driciru, & Rwetsiba, 2010).

During the 1960s, QENP had a vibrant wildlife research programme at Mweya, the Nuffield Institute of Tropical Animal Ecology (NUTAE), later becoming the Uganda Institute of Ecology (UIE). At this time, the elephant population varied from 1800-3500 (with migration between QENP and Virunga), whilst buffalo were fairly stable at 16000-19000 (Eltringham, 1977; Eltringham & Woodford, 1973; Field, 1971; Field & Laws, 1970). QEPA’s Uganda kob population, spread across open plain and around breeding leks, was estimated by ground and aerial counts at some 8000-12000 (Eltringham, 1973; Eltringham & Din, 1977; Modha & Eltringham, 1976).

During the political chaos of the Amin regime of the 1970s QEPA’s wildlife populations were hunted relentlessly by heavily-armed militias, with elephants targeted for ivory. By 1980 elephants and buffaloes in QEPA had been reduced by 80% to just 150 and 4200 respectively (Douglas-Hamilton et al., 1980; Eltringham & Malpas, 1980; Malpas, 1981), both by hunting and emigration to PNV. With peace restored in the mid-1980s, intensive conservation efforts by the (then) Uganda National Parks - now the Uganda Wildlife Authority (UWA) - and its development partners resulted in a steady recovery of wildlife populations. In 1995, some 16000 buffalos were recorded in their former ranges of grasslands and waterholes, whilst an estimated 31000 kob were mapped to former long-established ‘leks’, or breeding grounds (Lamprey & Michelmore, 1995).

Aerial counts of 2006, 2010 and 2014 in the landscape also included PNV in the Democratic Republic of Congo to obtain a more holistic view of wildlife trends across the Greater Virunga Landscape (Plumptre, Kujirakwinja, et al., 2010; Plumptre, Kujirakwinja, Owiunji, et al., 2007; Wanyama et al., 2014). These counts recorded that whilst QEPA’s immediate post-millennium recovery was holding firm, Virunga’s large mammal population declined by 90%, largely due to massive poaching during DRCs political instability of the early 2000s. Elephants all but disappeared from PNV, the survivors emigrating to QEPA (Keigwin, Wabukawo, Wasser, & Chapman, 2016). By 2010, the elephant population of QEPA had grown to 2 500-3 000 (Wanyama et al., 2014).

Recent advances in genetic sequencing have drawn attention to QEPA as the primary interface between the African savannah elephant *Loxodonta africana* and the African forest elephant of central Africa, *Loxodonta cyclotis* (Mondol et al., 2015), recently recognized as a separate species (Hart et al., 2021). The high degree of hybridization (>50%) of these two species in QEPA, occurring over multiple generations, suggests that pulses of immigration and genetic exchange have occurred over many decades.

In QEPA, buffalo and kob populations appeared stable during the early 2000’s (Plumptre, Kujirakwinja, Treves, et al., 2007). However, a sample-count conducted in 2010 indicated another major decline of these species by 50%, to just 8128 ± 3218 (95% c.l) and 8483 ±3486 (95% c.l) respectively (Plumptre, Kujirakwinja, et al., 2010). This decline has been attributed to renewed poaching and an outbreak of anthrax (Driciru et al., 2018). A contributing factor has been the spread of the unpalatable invasive grass species *Imperata cylindrica* (‘Cogon grass’) which has rendered many areas unsuitable for kob and buffalo (Ayebare, Kirunda, Nyago, & Nampindo, 2020; Jaksic-Born, 2009; Lock, 1985; Plumptre, Kirunda, et al., 2010). Against this background of increasing threats, the results of the most recent SRF-RSO survey of 2014 were encouraging, with 15 771 buffalo and 12 987 kob estimated (Wanyama et al., 2014). This survey suggested that once again the large mammal populations of QEPA were recovering rapidly.

To better understand these recent trends, the UWA and its NGO partner the Uganda Conservation Foundation (UCF) proposed that a high intensity aerial count be carried out to clarify this recovery and provide an updated baseline. This was commissioned as a sample count survey, as conducted in QEPA and across Africa, implemented according to the systematic reconnaissance flight (SRF) protocols where the sample unit is the strip-transect and the analysis is by the Jolly II method (Caughley, 1977; Gasaway, DuBois, Reed, & Harbo, 1986; Jachmann, 2001; Jolly, 1969; Norton-Griffiths, 1978). In the ‘traditional’ SRF, rear-seat-observers (RSOs) count animals to left and right of the aircraft within sample strips defined by markers on the aircraft. Where animals occur at low density or in spatially uneven or clumped distributions, population estimates derived from SRF sample counts often have high variance since there is a high probability of recording sample units with low-or zero-counts (Ferreira & Van Arde, 2009; Jachmann, 2001; Redfern, Viljoen, Kruger, & Getz, 2001).

In addition to such sampling errors, there are a range of factors that cause sample bias, including vegetation cover, animal size and colour, group size and the counting competency of crew (Caughley, 1974; Graham & Bell, 1989; Jachmann, 2002; Jacques, Jenks, Grovenburg, Klaver, & Deperno, 2014; Lee & Bond, 2016; Pollock & Kendall, 1987; Redfern et al., 2001; Tracey, Fleming, & Melville, 2008). RSO counting ability is a key factor, and recent SRF-RSO evaluation methods that re-analyse raw survey data (Craig, 2012; PAEAS, 2014) point to circumstances where RSOs have counted significantly different numbers of animals (Schlossberg, Chase, & Griffin, 2016), or where, per unit of counting effort, two separate survey aircraft flying interleaved transect patterns generate significantly different population estimates (Wanyama et al., 2014). This variability may mask significant changes in the wildlife population and the factors that cause them (Reilly, van Hensbergen, Eiselen, & Fleming, 2017).

QEPA’s wildlife is heavily clustered, with large herds of buffalo congregating in localized ranges around waterholes and kob concentrated in breeding grounds or ‘leks’ (Balmford, 1992; Deutsch, 1994); in such conditions, traditional SRF sampling will inevitably result in low precision (Jachmann, 2001). SRF counts of QEPA from the 1960s to the early 2000s were conducted at 6-8% sample intensity (Douglas-Hamilton et al., 1980; Eltringham & Din, 1977; Lamprey, 2000; Lamprey & Michelmore, 1995), increasing to 10-12% intensity in surveys since 2004 (Plumptre, Kujirakwinja, et al., 2010; Plumptre, Kujirakwinja, Owiunji, et al., 2007; Wanyama et al., 2014). Even with the increased sample intensity of recent surveys, where transects are flown with separation of 2.5 km, the ‘relative margin of error’ (RME), expressed as the ratio of 95% confidence limits to the estimate (CITES-MIKE, 2019; Norton-Griffiths, 1978), exceeds 50%, indicating low precision; for example the elephant estimate for Q.ENP of 2010 is calculated at 2502 ± 1441 (95% c.l), indicating an RME of 58%.

Our QEPA count is a multi-species count, but with particular emphasis on elephants. Given the high variance of previous counts, we explored the sampling intensity required to determine a trend (an effect) of a 40% change of the elephant population. We aimed to achieve an RME for elephants of ≤ 20%, widely seen as the ‘target precision’ for sample counts (CITES-MIKE, 2019; Ferreira & Van Arde, 2009; Norton-Griffiths, 1978). If sampling intensity can be increased, and variance between sampling units kept low, this 20% RME could, in theory, permit the significant detection of a 40% change in population, with the probability of a Type I error at *a of* 0.05 or, better, a Type II error at *β*= 0.2; ie the power of the test is ≥ 80% (Craig, 2012; Gasaway et al., 1986; Gerrodette, 1987; Reilly et al., 2017; Steidl, Hayes, & Schauber, 1997). In practice, an RME ≤ 20% is achievable for counts at a regional level where estimates for the component counting blocks or strata are combined (Chase et al., 2016). Here, the strata / block population estimates are additive, whilst the overall variance is the square root of the pooled stratum variances (Gasaway et al., 1986; Norton-Griffiths, 1978). Thus, the improvement of precision with increasing sample size is not linear but decreases in a diminishing return between survey cost and required precision.

An RME of ≤ 20% is very rarely achieved for a single stratum, which might be a survey block of around 1000-2000 km^2^, ie a main block of a small national park such as QEPA. However, despite this smaller size, the area might be a critical elephant refuge, and managers still need to determine if a population change is ‘significant’. Our approach was to improve RME to a target of 20% by implementing a sample intensity of 14%, with the possibility of doubling this to 28%. Due to the cost and practicalities of aerial sampling with the prescribed strip-width of 150 m (CITES-MIKE, 2019), sample sizes higher than 28% can only be simulated, and we derived models from the elephant group-size distribution recorded on the actual transects (Ferreira & Van Arde, 2009).

Meanwhile, to address detection bias, we used the ‘oblique camera count’ (OCC) survey method where the entire sample strip is captured using high-resolution digital cameras in order to obtain verifiable animal counts (Lamprey et al., 2020, 2019). This method uses standard CITES-MIKE RSO-survey parameters of strip width, but with the feature that imagery of the strip can be visually or electronically scanned, for minutes or even hours, to count animals; this is in contrast to the 5-7 seconds that an RSO has to visually detect and count animals in the sample strip during flight. The OCC method also provides a foundation for machine learning of wildlife by computer systems to improve counting efficiency (Delplanque, Foucher, Lejeune, Linchant, & Théau, 2021; Eikelboom et al., 2019).

RSO- and OCC-based counting methods are therefore significant different, and due caution should be applied to determining recent wildlife trends from surveys based on these two methods. Recognizing this, we aim to establish new baselines for QEPA, where every animal in the sample strip is indexed, verified and georeferenced, with the potential for third-party re-analysis. For future trends, the same image-based methods can be used, without the issue of variability in RSO counting ability. This paper presents the methods and the results of this OCC-based high-intensity survey and makes recommendations for the design of future surveys.

## 2. METHODS

### 2.1 Study area

The study area is the Queen Elizabeth Protected Area (QEPA), which comprises *Themeda triandra* and *Imperata cylindrica* grasslands, open woodlands with *Acacia sieberiana* and *A. gerrardii*, and thickets along the main channels and lakeshores with high emergent *Euphorbia candelabrum* and thorny *Capparis tomentosa* shrubs. The park is bisected in the centre by the dense Maramagambo Forest, which, together with the contiguous Kalinzu FR outside QEPA’s boundaries, covers 730 km^2^. Maramagambo-Kalinzu FRs are a surviving eastern extension of central African lowland forests (Aleman, Jarzyna, & Staver, 2018), dominated by *Parinari excelsa, Cynometra alexandrei* (‘ironwood’), *Warburgia ugandensis* and *Maesopsis eminii* (Langdale-Brown, Osmaston, & Wilson, 1964; Lock, 1985). Rainfall is approximately 700-900 mm per annum, received bimodally as ‘long rains’ of March-May and ‘short rains’ of September-November (Diem, Konecky, Salerno, & Hartter, 2019).

Figure 1 shows the census zone (CZ) of 2650 km^2^, which includes Queen Elizabeth National Park and the contiguous Kyambura and Kigezi Wildlife Reserves. As is usual in QEPA, the survey excludes the Maramagambo Forest, where forest cover is too dense for aerial detection of animals. The flight transects extended 2 km out from lake shorelines to capture local fishing activities. The analyses were conducted according to the three standard survey strata for QEPA, as applied in most previous SRF surveys, see Figure 1. Specifically, these are Stratum 1, ‘Ishasha Block’ (525 km^2^), from Kisenyi village south to Ishasha customs post; Stratum 2, ‘North Block’ (1288 km^2^), Kisenyi village north to Lake George; Stratum 3, ‘Dura Block’ (469 km^2^), the area north of Lake George.

**Figure 1.**
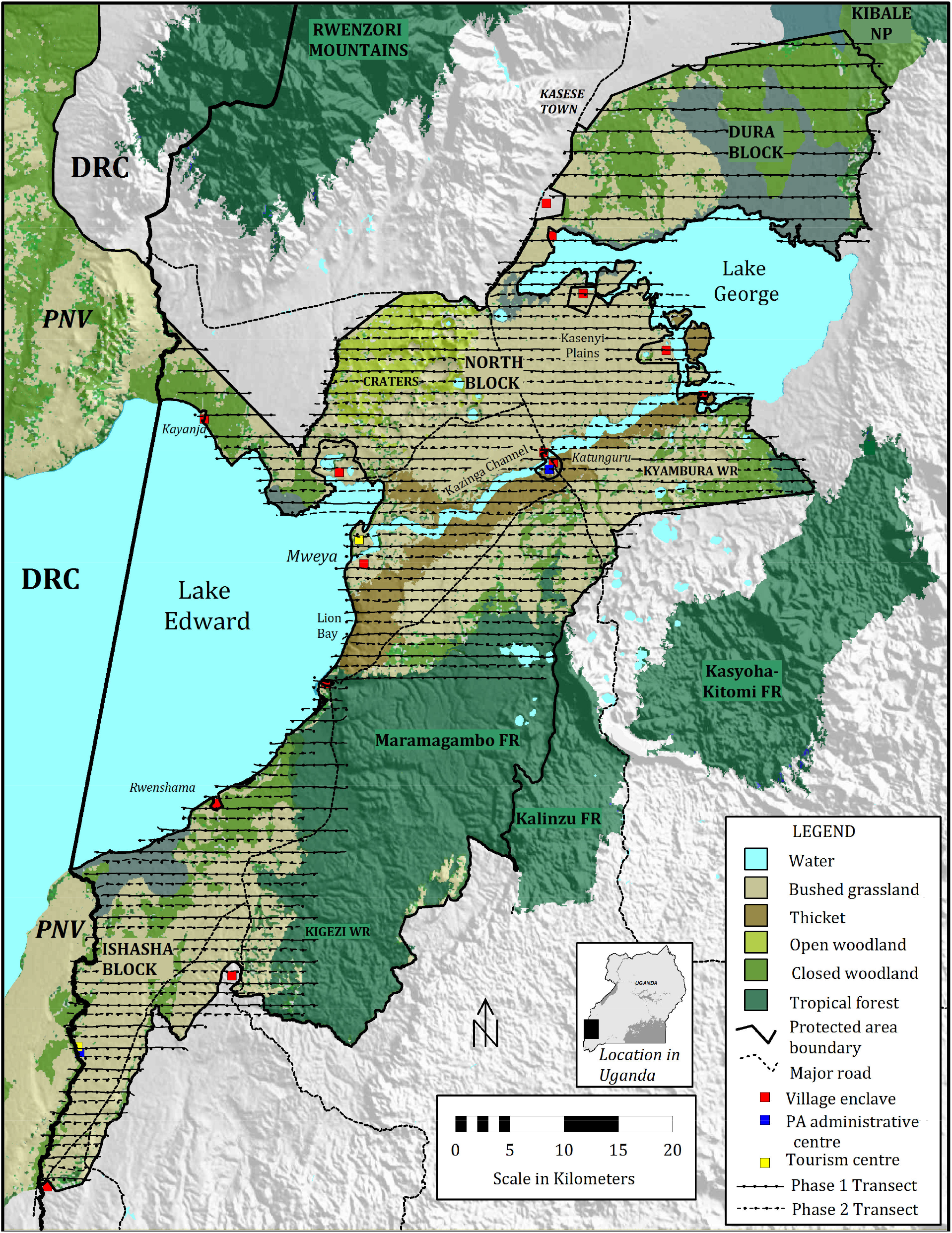
Map of the Queen Elizabeth Protected Area survey area and the location in south-western Uganda adjacent to Parc National des Virungas (PNV) in Democratic Republic of Congo (DRC). The map shows the three counting blocks of Ishasha, North and Dura, and the 2018 flight transects.

### 2.2 Survey design and navigation

QEPA has two elephant counting strata, North and Ishasha; elephants are generally absent from the third stratum Dura, north of Lake George. We flew transects in two phases; firstly at 2 km interval, giving a sample size of 14% (Phase 1); and then by flying ‘infill transects’ to achieve a spacing of 1 km, which increased the sample size to 28% to improve precision (Phase 2). A sample size of 28%-30% is essentially the highest practical trade-off between sampling effort and cost whilst adhering to 150 m sample strip-widths advocated in current elephant aerial survey standards (CITES-MIKE, 2019). This design accords with, and exceeds, the recommendation under CITES-MIKE (2019) for a sampling intensity of 20% for areas of size 1000-5000 km^2^, and resulted in a significantly narrower transect separation than the ‘tightest’ spacing of 2.5 km prescribed in the CITES-MIKE guidelines.

The survey was designed with parallel east-west flight-lines aligned to a Universal Transverse Mercator grid across the census zone. Phase 1 implemented a 2 km transect separation and was flown on 29-30 September 2018. Phase 2, flown on 1-2 October 2018, implemented the ‘infill’ sampling between the Phase 1 transects, to provide overall a 1 km separation. The flightlines were created in a MapViewer GIS system, for uploading to the aircraft aviation GPS prior to flight. Flight navigation was conducted using Garmin 96 and 196 aviation GPS units preloaded with the transects. During the survey, the tracklog was set to record at 4 s intervals for a full record of the flightlines, as well as providing the height-above-sea-level (HASL) for each tracklog point which forms the basis for post-processing of the actual HAGL, for each photopoint, using the SRTM DEM approach. The actual transects, as drawn from the GPS tracklog, are shown in Figure 1.

### 2.3 Aircraft and camera parameters

A Cessna 182 aircraft was used, registration 5Y-ATS, with large camera port in the floor for both Nikon cameras. The aircraft, operated without observers, was piloted by the lead author (RHL) with experience in SRF flying. Two 24-megapixel Nikon cameras (D5300, D3300) with 18-70 mm zoom lenses were angled to acquire continuous-strip imagery at 90° to the direction of travel, and at 45° inclination to vertical, see Figure 2 (a). The lenses were adjusted to 40 mm focal length, to nominally provide a strip width each side of 150 m, with groundsampling-distance (GSD) of 3 cm (Neumann, 2008; O’Connor, Smith, & James, 2017); calibrations were then tested in the strip-width calibration exercise. The cameras were mounted in dense vibration-absorbing foam blocks, which were sculpted to cradle each camera to achieve the required angle, see Figure 2(b).

**Figure 2.**
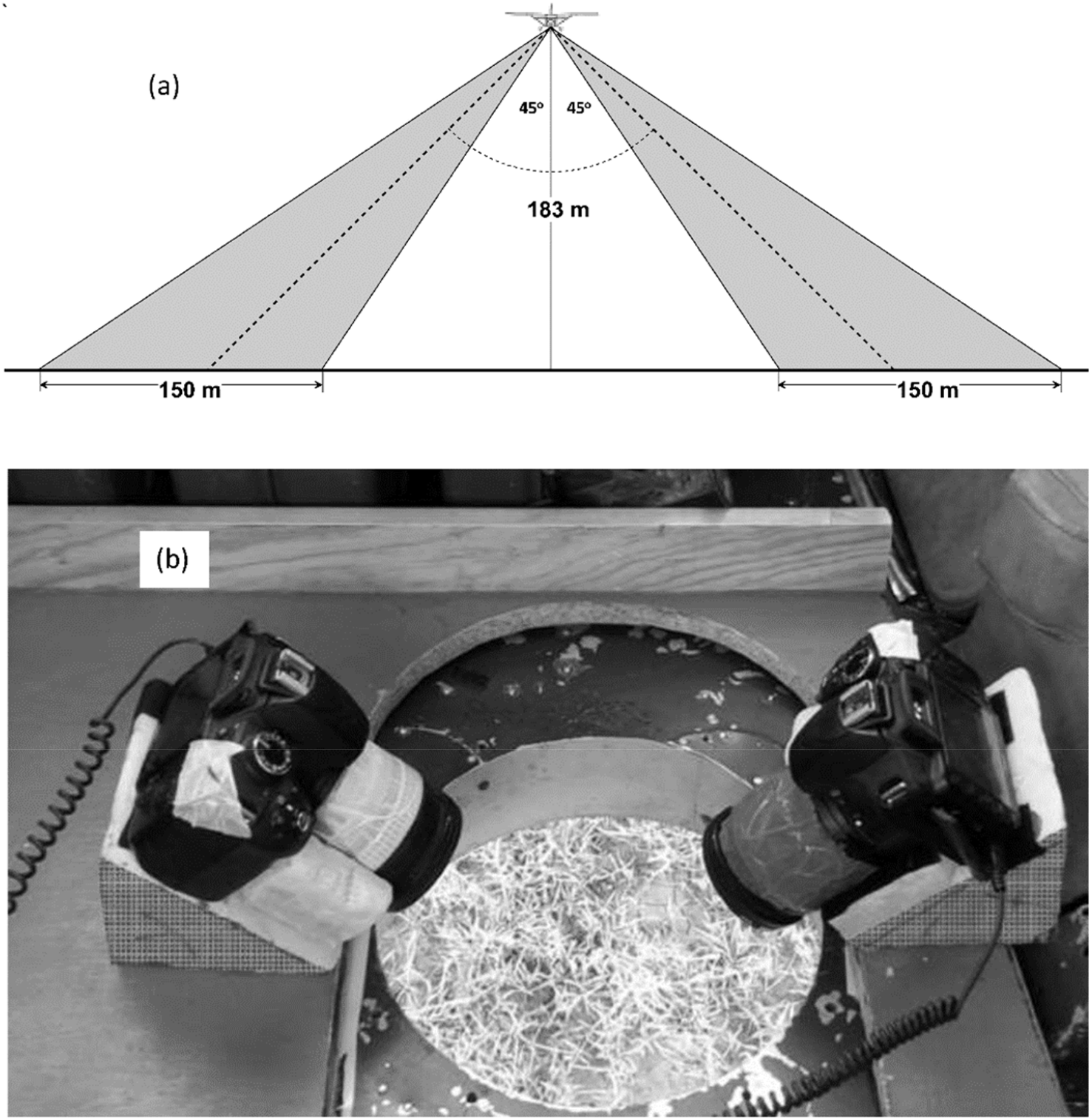
(a) Aircraft height and camera angles to obtain nominal strip width of 150m each side of the aircraft, or 300 m strip width in total. In practice the mean combined strip width obtained was 288 m, as measured from the GD-HAGL and strip-width calibration. The diagram is drawn to scale (except aircraft), (b) Cameras were installed in simple vibration-absorbing aeronautical foam blocks and angled for clear field-of-view through the aircraft camera hatch. Critical camera settings such as focus and aperture were taped to avoid disturbance.

The cameras were triggered by intervalometers at 2 s interval, corresponding to 113 m of forward movement of the aircraft at aircraft ground-speed of 110 knots (204 km/h); in practice we selected 105 knots (194 km/h) as the target speed. Camera clocks were synchronised to the aviation GPS clock for subsequent geo-location of each image. Approximately 24000 images were acquired for the main 2 km transects, and 19000 images for the infill transects. Geo-location was conducted for the full airphoto set, using RoboGeo. This procedure, taking several days of processing time generated accurate locations (50-100 m) for each photo that were then used to generate the GIS shapefile. In addition, the procedure generated the height-above-sea-level for each image, for subsequent processing over the SRTM DEM model, to generate HAGL for each photo.

### 2.4 Height-above-ground and strip-width calibration

The HAGL is the primary factor in determining sample strip width, and must be regulated within certain practical limits; the standard is 300-350 ft a.g.l with RSOs (PAEAS, 2014), or 500 - 600 ft with camera systems (Lamprey et al., 2020). Generally, HAGL is controlled by reference to radar-or laser-altimeters. However, these are expensive, difficult to install and often unreliable. Hence for the Q.ENP OCC survey the GPS-DEM approach was used, whereby height-above-ground is indicated as the separation of the aircraft GPS from the elevation of the ground below, using the Shuttle RadarTopographic Mission (SRTM) elevation model (Lamprey et al., 2020). For QENP, the prescribed HAGL was set at 600 ft above ground level (a.g.l), and prescribed height-to-fly to achieve this separation is indicated on the GPS at 1 km intervals (see Figure 1).

Strip-width-calibration (SWC) provides the calibration of HAGL to RSO-based or OCC strip width for each imaging strip. The strip for the QEPA survey was nominally prescribed at 150 m each side of the aircraft, in accordance with standard SRF practice with rear-seat observers (CITES-MIKE, 2019; PAEAS, 2014). The SWC was conducted by photographic overpasses at height increments of 100 ft (300 ft up to 1100 ft a.g.l) over the Mweya airfield. Markers were placed alongside the runway at 30 m intervals, with finer definition derived from a perspective grid superimposed on each overpass image. Figure 3(a) shows the SWC and frame footprint calibration for the QEPA survey, the latter derived for ‘point sample’ analysis of alternate images. The calibration was slightly different for left and right cameras due to extremely sensitive settings on the zoom lenses, but in practice, and as with RSO-based surveys, the two strips are combined to form the ‘strip-transect’. At 600 ft HAGL, the combined strip width is 282 m. Figure 3(b) shows a histogram of HAGL during the flights, measured using the GPS-DEM approach as the difference of image height with the underlying terrain at that precise point. The mean height obtained is 634 ft ± 73 (SD), indicating satisfactory height control from the high Craters in the west, across the central plains to undulating terrain of the Maramagambo forest and Albertine Rift (‘Kichwamba’) escarpment in the east.

**Figure 3.**
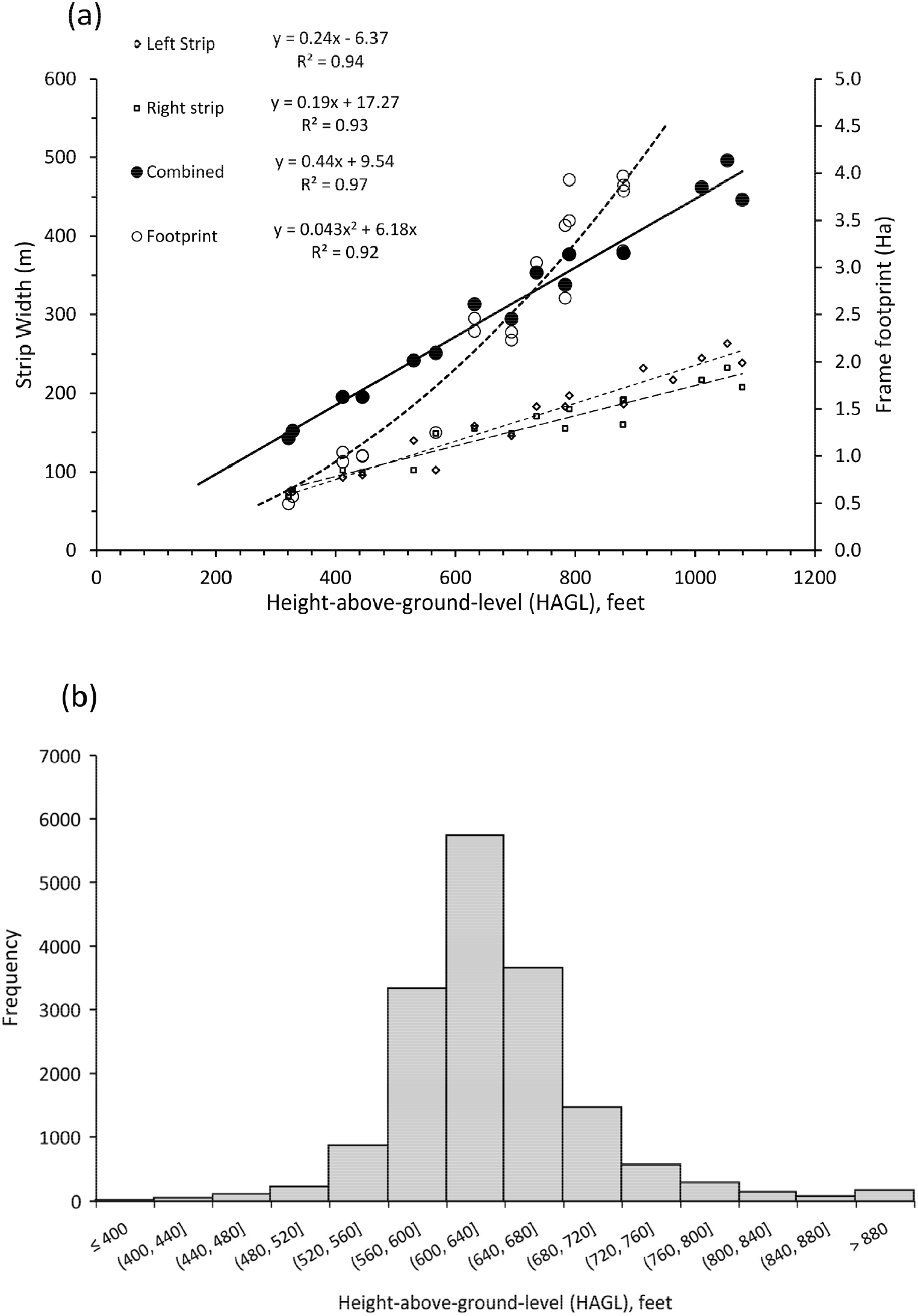
(a) The strip-width-calibration and frame footprint calibration for the QEPA survey. Note, for clarity of the footprint regression equation, the area unit is m^2^, although the scale (right Y-axis) is in hectares, (b) Histogram of HAGL during the flights, measured from even-code images where N=16751, mean = 634, SD =73.0. This is measured using the GPS-DEM method and is calculated as the difference of image height (the GPS elevation) with the underlying SRTM terrain elevation at that precise point. HAGLs higher than 800 ft are derived from climb/descents from the eastern Albertine escarpment.

### 2.5 Image interpretation

From the image analysis, we counted 13 wildlife species, 27 anthropogenic variables and 8 landscape/ vegetation classes, these being water, wetland, grassland, *Imperata cylindrica* grassland, wooded grassland, thicket and forest. *Imperata* is a tussock grass highly visible in airphotos, especially if it has not been recently burned. Estimation of vegetation variables is simply the proportion of aerial images in the subunit (usually 18 images per 1 km subunit) where the class is recorded as the dominant type in the image.

The digital airphotos were interpreted by a team of four Ugandan interpreters, originally trained in similar exercises in Uganda and Kenya (Lamprey et al., 2020, 2019). The interpreters worked as two teams, A and B, with a systematic alternating allocation of images by transect and camera side (Lamprey et al., 2020) to ensure that interpretation was unbiased with respect to high-density wildlife areas, or to camera quality. Initially, the images were renamed to a standard protocol that includes dates, camera side, and the folders and frame numbers as preset by the Nikon cameras. Images were then projected using standard Microsoft Windows Photo Viewer (for visual panning/ scanning) and IrfanView for photo enhancement and photo EXIF information. For each frame, data on species numbers were entered into a standard data spread sheet by the fifth member of the team, the data entry clerk.

Where a large herd (eg elephants, kob) spans overlapping photos, animals in the overlap area are counted into the even-number photo. Thus, every photo coded by the camera with an even-number is a total count (evenframe-summation or ‘EFS’), whilst every odd-number photo ‘filled’ the gaps between, see Figure 4. This approach provides a dataset of even-code photos where there is no possibility of double counting. Figure 5(a) shows an image of buffalo and Uganda kob, with enlargements to show detail, and Figure 5(b) shows enlargement of receding overlap frames, at the outer edge of the strip, to reveal hidden elephants under trees.

**Figure 4.**
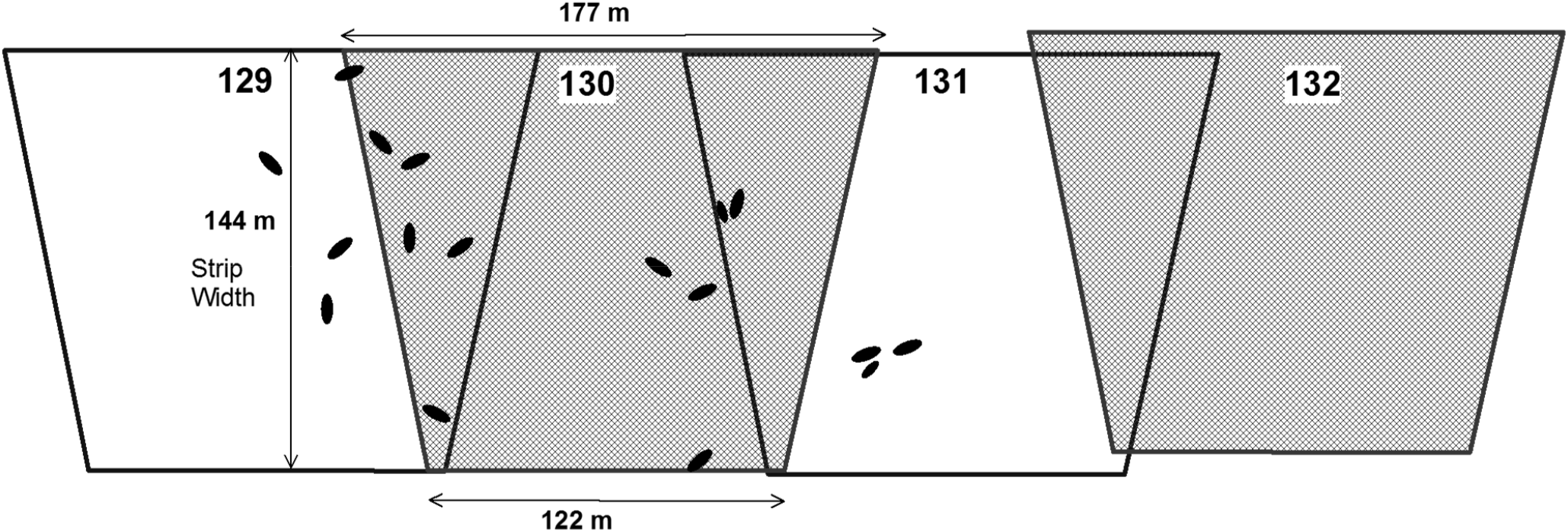
Diagrammatic image footprint string, left side camera, drawn to scale. Frame numbers are given at top of each image. In the alternate image ‘point-sample’ assignment (by even-frame-numbers), the animals shown as black dots to left are assigned as 3 animals to image 129,11 to 130, 3 to 131.

**Figure 5.**
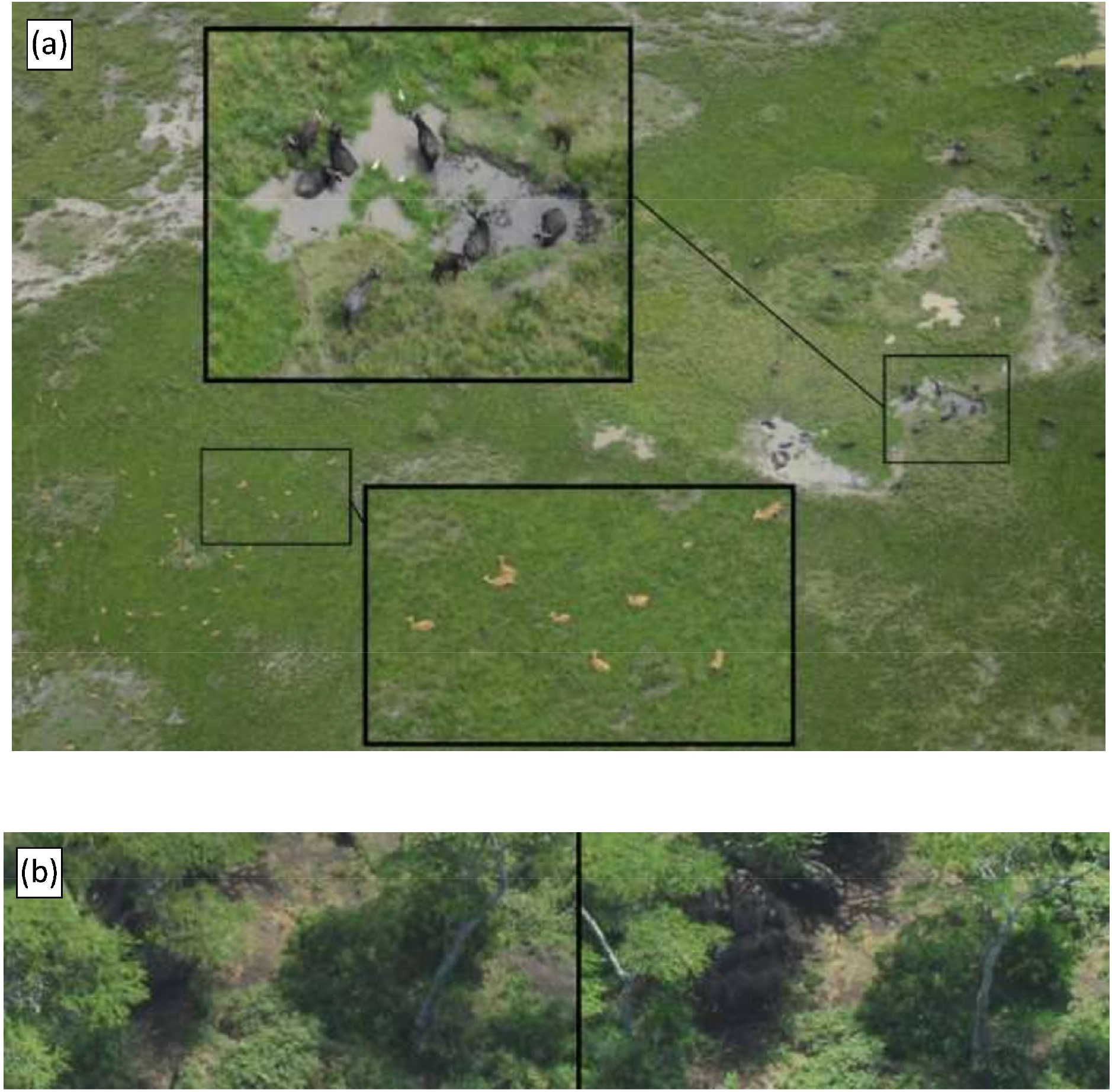
(a) Image of buffalo and Uganda kob in Ishasha Block, with enlargements to show detail, (b) Elephants in Craters area becoming visible under trees (aircraft moving rightwards) in the overlap with the receding frame (left) along the transect.

Initially only the Phase 1 images, constituting a 14% sample intensity, were analyzed for the Report to the survey sponsors. However, it was soon realized that although the population estimate for elephants in Phase 1 was higher than expected (3953 ± 1714,95% c.l), it lacked precision with an RME of 43%. We therefore interpreted the additional 19000 images of Phase 2 to increase the sample size from 14% to 28%, and thereby improve precision. Image analysis took place over a six-week period in October-November 2018, with a total of 37 000 photos interpreted.

### 2.6 Data analysis

Detection biases might arise from differences in counting abilities between interpreters, differences in lighting conditions of the two phases of the count (main transects afternoon, infill transects morning) or quality of imagery of the left and right cameras. To test for counting biases between Team A and Team B, we followed the guideline set down by the Pan-African Elephant Aerial Census (PAEAS, 2014), where the chi-square (χ^2^) test is prescribed for comparison of species occurrences and numbers between the teams. Similarly, we used chisquare to test for differences between the left and right camera systems.

The survey data were then analyzed according to the standard Jolly II method for unequal sized sample units (Caughley, 1977; Jachmann, 2001; Jolly, 1969; Norton-Griffiths, 1978). If we consider one species at a time, then the population estimate is derived from a simple ratio estimator (Cochran, 1977) for the average density of the species estimated across all transects:

*z* = the area sampled for each transect (the sample unit)

*Z* = the total area under survey

*y* = the number of animals enumerated along the transect

*R* = total number of animals counted in all transects / total area sampled (ie the average density) = ∑*y/∑z*

The population estimate is then found by:

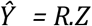

and the precision of the population estimate is calculated as:

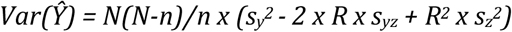

Where *S*y^2^ = the variance between the numbers in all transects; *S*_z_^2^ = the variance between the areas of all transects; and *S*_yz_ = the covariance between numbers and the area of the transects. The standard error is calculated as 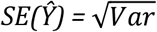, where *n* = the number of sample units in the sample, and *N* = the total number of sample units from which the sample was drawn. The 95% confidence limits (95% c.l) of a survey estimate are calculated by multiplying the standard error (SE) by the *t* value for n-1 degrees of freedom (where *t* approximates to 1.96 for high values of n).

Our analysis for QEPA is based on the standard Jolly II sample-size estimation where transect sampled area (km^2^) = length (km) x mean strip width (km). In the OCC method, use of the length x width metric assumes that interpreters have been able to identify and eliminate animals in image overlaps in the EFS process. To determine how effective they were in eliminating double-imaged groups, we conducted a further Jolly II analysis based on counts from the only the ‘point-samples’ of alternate EFS images, where all animals are counted. Here, the transect sample area *∑z* (sq.km) = *∑ai* where *i* is every alternate image 2,4, 6.. and where *a* is the image ‘footprint’ area (km^2^) of the *i*th image. Population estimates were then compared between continuous strip and point samples.

### 2.7 Sample intensity and precision

For future sample survey design in QEPA, we test the effects of both reducing and intensifying sample intensity on precision using North Block. For deselection, to test sample intensities below 28%, we draw every 4^th^ transect to derive a 4 km spacing with an 8% sample with 4 possible allocations; each of these is subject to a Jolly II analysis. We also draw every 2^nd^ transect to derive a 2 km spacing with an 15% sample with 2 possible allocations. Sample intensities higher than the existing 28% sample intensity must rely on simulation. Here we create and model the frequency distribution of elephant group sizes in North Block detected in the imagery, and then randomize from that distribution to establish herds along additional simulated transects up to a sample size of 100%. Elephants in North Block are not randomly scattered across the stratum, but are distinctively congregated along the Kazinga Channel, with smaller clusters in the Craters and towards Maramagambo. Therefore, our distribution of simulated elephants is by random positioning along additional transects to three ‘sub-strata’ (Craters, Kazinga, East), depending on their densities within these strata in the original (unsimulated) dataset. We run Jolly II on all sample intensities to determine the effect of sample size on RME in QEPA’s North Block.

In a second test of elephant estimate RME, we calculate the 95% confidence limits of the Jolly II estimate by an entirely different method, bootstrapping. Here we recalculate the Jolly II estimate *Ŷ = R*.*Z* through 1000 bootstrap iterations of random resampling of transects with replacement. The resulting distribution of Jolly II estimates is near-normal (Shapiro-Wilk test, W=0.999, p = 0.971, n=1000), and confidence limits are then determined by the percentile method, as lying between the 50^th^ and 950^th^ observations of the distribution (Diciccio & Romano, 1988). As before, the transect dataset can be de-sampled to provide lower sampling intensities.

We test a third approach, the experimental ‘Marriott Method’ to derive the variance and confidence limits of the population estimate (Marriott & Wint, 1985; Wint, 1996). Instead of using the single dimension of transect direction, perpendicular to a ‘baseline’, for the measure of variance, the Marriott Method uses the transect subunit as the sampling unit; variance is determined in two dimensions from the unit. The variance of the estimate Ŷ is calculated as:

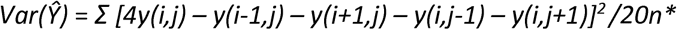

where i,j = the array coordinates (eg row, column) of the grid cell; *y* = the count of animals in the grid cell (equivalent to *y* in Jolly II); n = number of cells with four surrounding neighbours; n* = the number of sample units (subunits). The Marriott Method relies on Ŷ = *R*.*Z* for the population estimate, and therefore this estimate is the same as that derived from Jolly II. However, the precision is improved where animals are clustered in particular localities within strata, ie there are many zeros in the sample units. The method has been used extensively in Botswana for wildlife monitoring (BASIS, 2000), but this use in QEPA is the first test of the procedure in East Africa. We test the method for two species, elephants and Uganda kob. A similar experimental approach of adaptive sampling of clusters is described by Khaemba & Stein (2002) which shows promise but has not been operationalized in counts.

### 2.8 Determining power of the sample intensity to measure elephant population changes

We test the power of our sampling intensity in determining future trends in elephants in QEPA. As typical with power analyses we have the acceptable probability *α* of avoiding a Type I error if the survey concluded that change had occurred, when in fact it had not occurred (Gerrodette, 1987; Steidl et al., 1997); *α* is commonly set at 0.05, but there are strong cases in sampling of biological populations to raising this to 0.10 (Gibbs, Droege, & Eagle, 1998). Secondly, we have the acceptable threshold of a Type II error, *β*, if we concluded that no change had taken place, when in fact it had; *βis* usually 0.2. The power of the test is 1-*β*.

*We* use the approach of Gasaway et al. (1986) to test the power in detecting a ‘consequential change of difference’ (CD), meaning the ‘effect’ or the percentage change of the elephant population that triggers management concern. Here, for a two-tailed test with α = 0.05, *t* and *p* are calculated in the normal way for comparison of two survey estimates, whilst the Type II probability *β* is calculated for *t°* at *vt* degrees of freedom, and variance *V* for estimates *Ŷ_1_* (elephant estimate of start survey) and *Ŷ*_*2*_ (elephant estimate of second comparative survey) respectively, where:

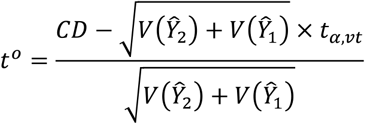

For our variance of *Ŷ_1_* and *Ŷ_2_*, *we* use the variance for our Jolly II estimate for a sampling intensity of 28%, assuming that the previous population estimate *Ŷ_1_* sampled at the same intensity would have a coefficient-of-variation-of-the-mean *(CV*_*mean*_ = *SE/Ŷ*) similar to *Ŷ_2_*, a not unreasonable assumption (Barnes, 2002; Gasaway et al., 1986).

What is an acceptable ‘consequential change of difference’? Supposing that we were to accept a 40% change (our ‘effect’) in elephant populations in QEPA as our trigger for concern, is our sample intensity sufficient to avoiding a Type I error at α = 0.05 or, better still, a Type II error at *β* = 0.2? We chose the previous RSO-based count in 2014 as an example of a start point, with 2913 elephants and increase the population (CD) by 3 increments, 40%, 60% and 80%. We apply this process at 6 different sampling intensities in the North Block simulation, 8%, 15%, 28%, 42%, 63% and 95%. We generate *t* for testing significance of the difference between the estimates *Ŷ*_1_ and *Ŷ*_2_, and compare the probability *p* at df (number of transects, both counts - 2) against α = 0.05. We calculate t° as above, and do the same for estimating *β*. Since *t* and *t*° vary according to *CD, V* and *df, we* repeat this estimation for CDs of 40%, 60% and 80%, measuring variance and the C*V* for each change of sample intensity. This analysis gives the level of sampling, ie how many transects we must fly across QEPA North Block with combined strip width 300 m, that are required to avoid a Type I error at α = 0.05 (or possibly 0.10) and a Type II error at *β* = 0.2, if we wanted to measure an effect size of 40%, 60% or 80% elephant population change since the last survey.

## 3. RESULTS

### 3.1 Wildlife population estimates

Table 1 shows the results of the Jolly II analysis for the three strata, using both the Phase 1 and Phase 2 images and sample intensity of 28%. The table shows the all-image analysis (where the sample is a continuous strip of overlapping frames), and of the EFS where the sample strip is comprised of separated independent ‘footprints’. There is no significant difference between all-image and EFS population estimates, and we retain the all-image estimates. Figure 6(a) presents the overall trend in elephants for QEPA since surveys began in the 1960s, together with 95% confidence limits of selected estimates, where data are available in the original study reports to derive these. Figure 6(b) shows the population estimates over time for buffalo and hippopotamus, and Figure 6(c) for Uganda kob, waterbuck and topi. These trends are discussed in detail below, for each species.

**Table 1.** Population estimates Ŷ (bold), standard errors (SE) and 95% confidence limits for the three survey blocks of Ishasha, North and Dura and for the QEPA totals. At right are the estimates Ŷ_EFS_ and standard errors SE_EFS_ derived from the even-frame summation (EFS), showing no significant difference with the all-frame analysis and therefore that double-counting in frame overlaps was not significant.

|  | Ishasha |  | North |  | Dura |  | QEPA Estimate |  | 95% Confidence Limits |  | QEPA Estimate-EFS |  |
| --- | --- | --- | --- | --- | --- | --- | --- | --- | --- | --- | --- | --- |
| | $\hat{Y}$ | SE | $\hat{Y}$ | SE | $\hat{Y}$ | SE | $\hat{Y}$ | SE | Lower | Upper | $\hat{Y}_{EFS}$ | $SE_{EFS}$ |
| **Elephant** | 839 | 194 | 3872 | 521 | 0 | 0 | **4711** | 556 | **3605** | **5817** | 4706 | 610 |
| **El. Carcass recent** | 0 | 0 | 4 | 3 | 0 | 0 | **4** | 3 | **0** | **10** | 5 | 5 |
| **El. Carcass old** | 14 | 7 | 43 | 10 | 0 | 0 | **58** | 13 | **32** | **84** | 62 | 19 |
| **Buffalo** | 5393 | 1563 | 8729 | 1170 | 494 | 289 | **14616** | 1974 | **10688** | **18544** | 14443 | 2565 |
| **Topi** | 1539 | 583 | 11 | 7 | 0 | 0 | **1550** | 583 | **390** | **2710** | 1726 | 720 |
| **Uganda kob** | 7184 | 1278 | 9290 | 1634 | 2140 | 925 | **18613** | 2271 | **14094** | **23132** | 18006 | 2402 |
| **Waterbuck** | 690 | 160 | 4697 | 510 | 439 | 230 | **5826** | 582 | **4668** | **6984** | 5685 | 644 |
| **Warthog** | 386 | 66 | 1983 | 248 | 82 | 46 | **2451** | 261 | **1932** | **2970** | 2264 | 275 |
| **Giant forest hog** | 42 | 22 | 290 | 79 | 14 | 13 | **346** | 83 | **181** | **511** | 299 | 90 |
| **Hippo** | 817 | 219 | 4734 | 522 | 165 | 147 | **5716** | 585 | **4552** | **6880** | 5645 | 732 |
| **Crocodile** | 7 | 4 | 127 | 49 | 0 | 0 | **134** | 49 | **36** | **232** | 159 | 58 |
| **Cattle** | 78 | 43 | 5374 | 1365 | 405 | 281 | **5857** | 1394 | **3083** | **8631** | 5469 | 1525 |
| **Goat** | 697 | 286 | 1571 | 390 | 48 | 48 | **2316** | 486 | **1349** | **3283** | 2155 | 477 |

**Figure 6.**
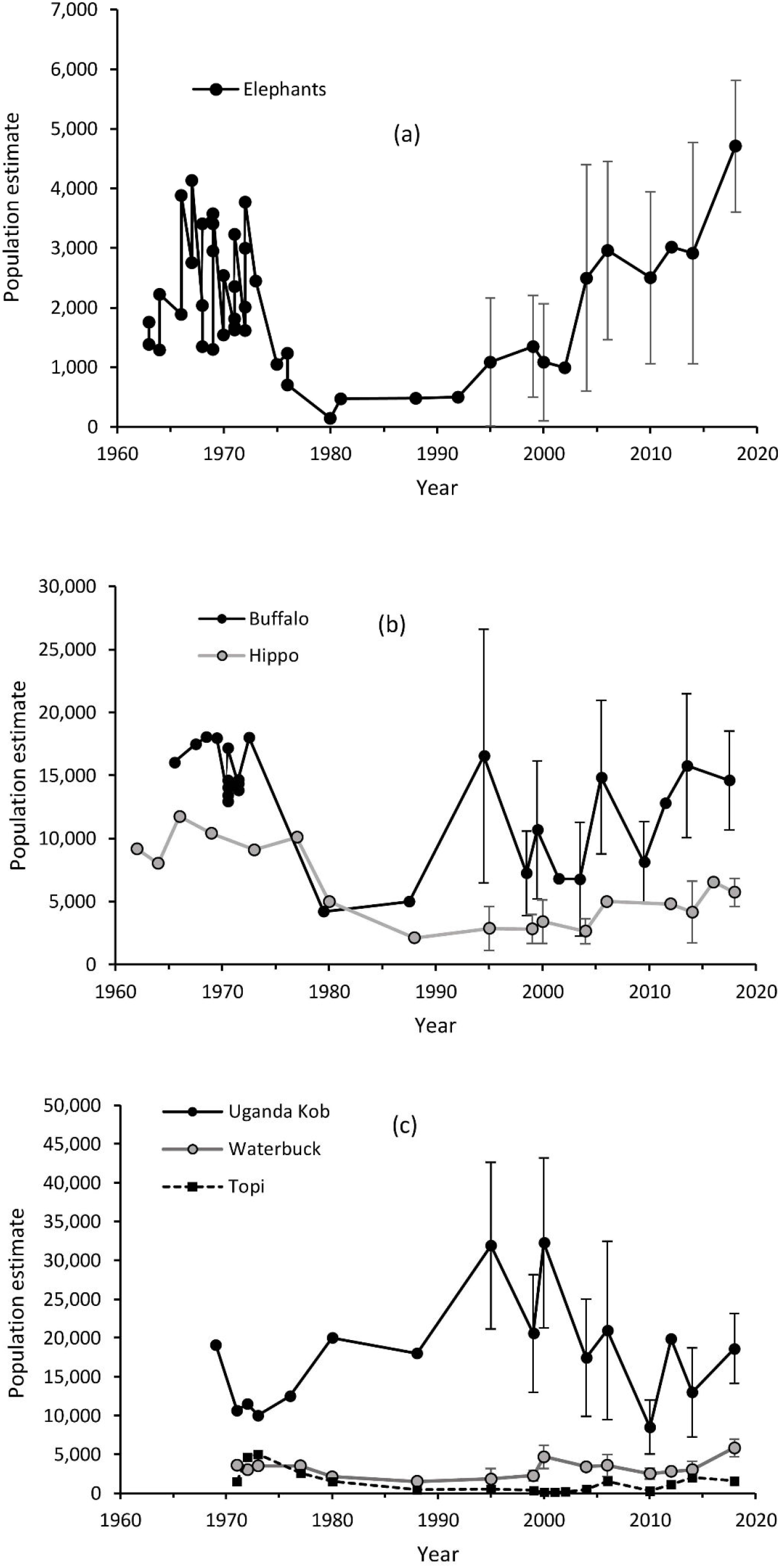
Graphs (a) to (c) show trends for a the main species of QEPA. Recent estimates from 1995 onwards derived from sample colunts show the 95% c.l of the estimates, where the standard errors are available in the literature.

In generating the population estimates shown in Table, 1, we tested for counting bias. Table 2 shows the results of chi-squared tests to determine possible detection or counting biases between the two interpreter teams, A and B, and for the left and right camera systems. Here, according to PAEAS elephant counting standards (PAEAS, 2014), we test for differences in the detection ‘occurrence’ rate for all species, regardless of herd size. We find no differences in interpreter detection for elephant, buffalo, Uganda kob, topi and hippo. However, significant differences exist for the more cryptic species of waterbuck with *χ*^2^ (1, N = 483) = 15.67, *p* < 0.001, warthog with *χ*^2^ (1, N = 246) = 13.67, *p* < 0.001 and giant forest hog *χ*^2^ (1, N = 39) = 9.26, *p* < 0.01, with Team B more effective at detecting these three species. Occurrence rates from the left and right camera were not significantly different; the exception was buffalo, with 162 occurrences on the left and 126 on the right, with *χ*^2^ (l, N = 288) = 4.62, *p* < 0.03; we do not consider this to be unusual, given the extremely clumped nature of buffalo herds. We conclude that for the large species there are no major biases in the interpreter counting. There are small biases in the dataset for smaller or cryptic species that lower the population estimates; these might be offset using corrections factors, but these are small and we do not implement them in this study.

**Table 2.** Results of *χ*^2^ tests to compare counts of Team and Team B, and from left and right side cameras.

|  | Compare Interpreters, Teams A and B |  |  |  | Compare Cameras, left & right |  |  |  |
| --- | --- | --- | --- | --- | --- | --- | --- | --- |
| | Team A | Team B | $\chi^2$ | $p$ | Left | Right | $\chi^2$ | $p$ |
| Total Images in Analysis | 18450 | 18427 |  |  | 18456 | 18422 |  |  |
| Elephant corrected for full | 136 | 147 | 0.57 | 0.45 | 137 | 146 | 0.42 | 0.52 |
| El. Carcass recent | 1 |  | 1.00 | 0.48 | 1 |  | 1.00 | 0.48 |
| El. Carcass old | 5 | 10 | 1.67 | 0.20 | 4 | 11 | 3.27 | 0.07 |
| Bones, White | 260 | 246 | 0.39 | 0.53 | 265 | 241 | 1.14 | 0.29 |
| Buffalo | 133 | 155 | 1.76 | 0.18 | 162 | 126 | 4.62 | 0.03 |
| Topi | 30 | 39 | 1.17 | 0.28 | 28 | 41 | 2.45 | 0.12 |
| Uganda kob | 396 | 435 | 1.83 | 0.18 | 398 | 433 | 1.48 | 0.22 |
| Waterbuck | 198 | 285 | 15.67 | 0.00 | 236 | 247 | 0.25 | 0.62 |
| Warthog | 94 | 152 | 13.67 | 0.00 | 124 | 122 | 0.02 | 0.90 |
| Giant forest hog | 10 | 29 | 9.26 | 0.00 | 20 | 19 | 0.03 | 0.87 |
| Hippo | 137 | 139 | 0.01 | 0.90 | 140 | 136 | 0.06 | 0.81 |
| Crocodile | 4 | 11 | 3.27 | 0.07 | 7 | 8 | 0.07 | 0.80 |
| Cattle | 33 | 36 | 0.13 | 0.72 | 38 | 31 | 0.71 | 0.40 |
| Roof (grass) | 221 | 270 | 4.89 | 0.03 | 282 | 209 | 10.85 | 0.00 |
| Roof (mabati) | 317 | 306 | 0.19 | 0.66 | 341 | 282 | 5.59 | 0.02 |

### 3.2 Elephant population and distribution

Inclusion of the Phase 2 imagery improved the RME precision of the QEPA elephant estimate from 43% to 23%. The elephant population is estimated at 4711 ± 1106 (95% c.l), the highest estimate in QEPA since surveys began in the 1950s and 60s, and the highest recorded precision for this species in QEPA. This elephant estimate is 62% higher than the previous 2014 RSO-based estimate of 2 913 elephants ± 1826 (95% c.l) (Wanyama et al., 2014). Figure 7 shows the elephant distribution, the highest densities being in the thicket areas along Kazinga Channel. The distribution maps of this and previous surveys suggest a recent eastwards movement towards the Kazinga confluence with Lake George, Akika Island and the northern areas of Kyambura WR. The elephant density in Maramagambo’s peripheral thickets and woodlands is calculated from our surveys at 3.3 elephants.km^−2^, one of the highest for any counting block in Africa. If such densities are characteristic within Maramagambo Forest itself, which was not surveyed, the elephant population of QEPA would be increased by a further 1200-1400 elephants.

**Figure 7.**
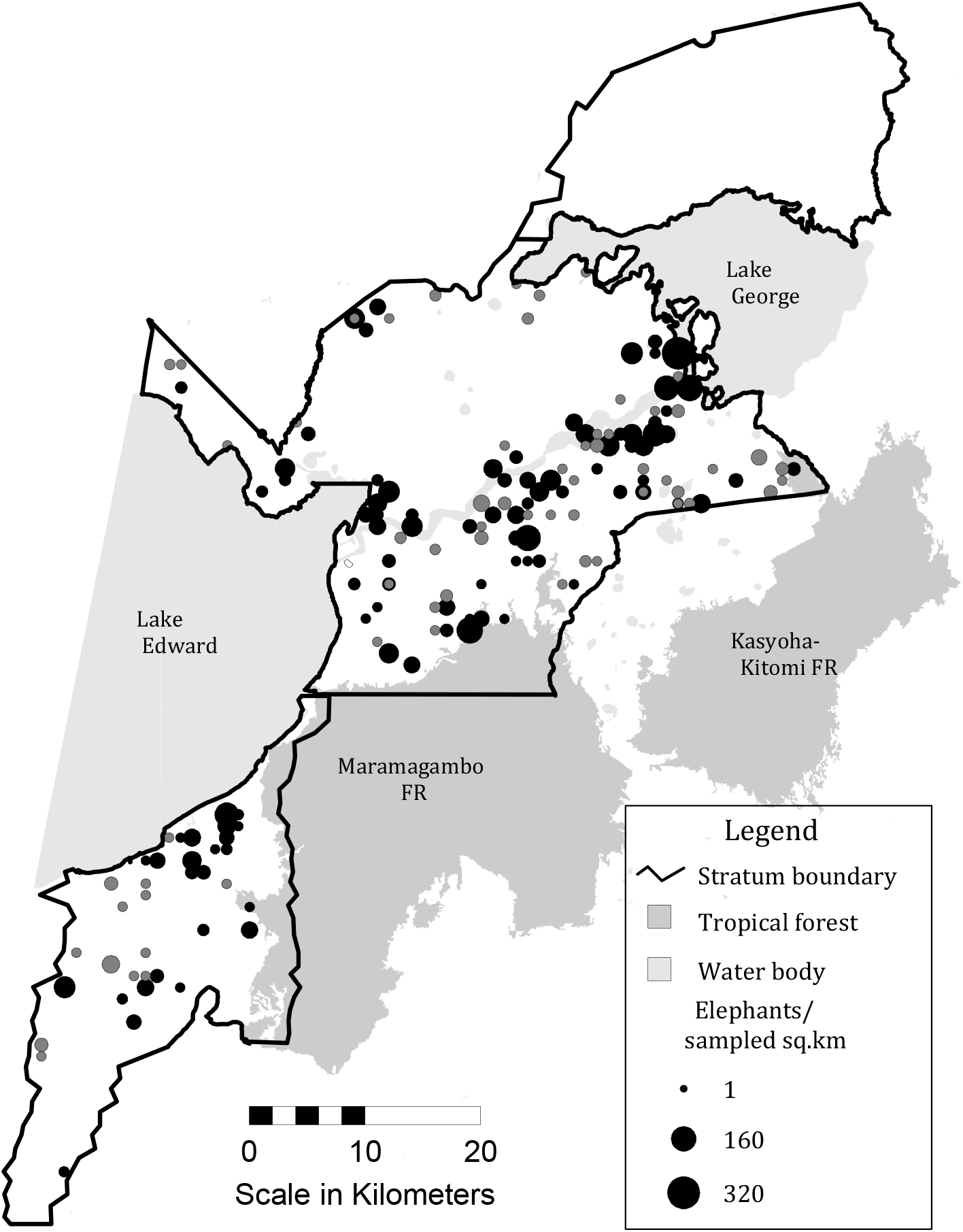
Distribution and density of elephants in QEPA. Black circles show the distribution of family herds and grey circles bull groups.

The elephant ‘carcass ratio’ (CR), where CR = carcasses/live elephants + carcasses, is an indicator of mortality rates in elephants (Douglas-Hamilton & Burrill, 1991). Generally, a CR of 3-7% indicates natural mortality, whilst a CR ≥ 10% suggests increased poaching and is cause for concern. ‘Rot-rate’ is a factor in persistence of carcasses; in the very humid conditions of QE elephant carcasses rot and disappear very quickly. In the Q.ENP 2018 survey, just eight elephant carcasses were counted, with one recent in the thickets adjacent to Kazinga Channel east of Katunguru. The overall estimate is 57 ± 19 (SE), giving a very low carcass ratio of just 1.4%. This reflects low mortality, and poaching is not significant. The carcass ratio in the GEC 2014 count was 3.1%, also low, whilst the 2010 count indicated only one carcass seen in QEPA.

### 3.3 Precision and accuracy of elephant estimates

Despite the magnitude of their differences, the estimates of 2913 elephants in 2014 and 4711 elephants in 2018 are not significantly different in a two-tailed test, where *t =* 1.675, df = 129, p = 0.096. This is largely due to the low sampling intensity of 12% (2.5 km transect spacing) in 2014, and the consequently high RME of 62%. This difference becomes significant at *α* = 0.05 if the test is one-tailed, ie the null hypothesis is that Ŷ_2018_ > Ŷ_2014_, and where *p*=0.048. Clearly, determining trends using the 2014 sample size of 12% (transect separation 2.5 km) is not powerful.

For future surveys, how far do we need to increase sampling intensity to indicate a significant increase in elephant population of QEPA? Figure 8 shows the effect of sampling intensity on elephant population estimates and RME for North Block, which contains over 80% of QEPA’s elephants. Excluding the small Kayanja arm of North Block which could not be sampled at 1 km interval due to weather conditions in Phase 2, the estimate is 3632 elephants ± 1056 (95% c.l.). Fig 8(a) shows the frequency distribution of elephant group size in OCC images, which is best modelled with a gamma or lognormal distribution; we chose the gamma with k-parameter = 0.828 and beta-parameter = 9.709; excluding a single large ‘outlier’ herd of 85 elephants, this gamma distribution returns *χ*^*2*^ = 19-6, df = 14, *p* = .13, indicating no significant difference between the model and the distribution. Randomization of this gamma-distribution enables a simulation of elephant herds along 3.4 parallel transects to the original with the same mean strip width (combined) of 291 m up to 1000 m, ie a 100% sample; these herds were distributed according to the original densities of the three sub-strata (Craters, Kazinga, East) of North Block.

**Figure 8.**
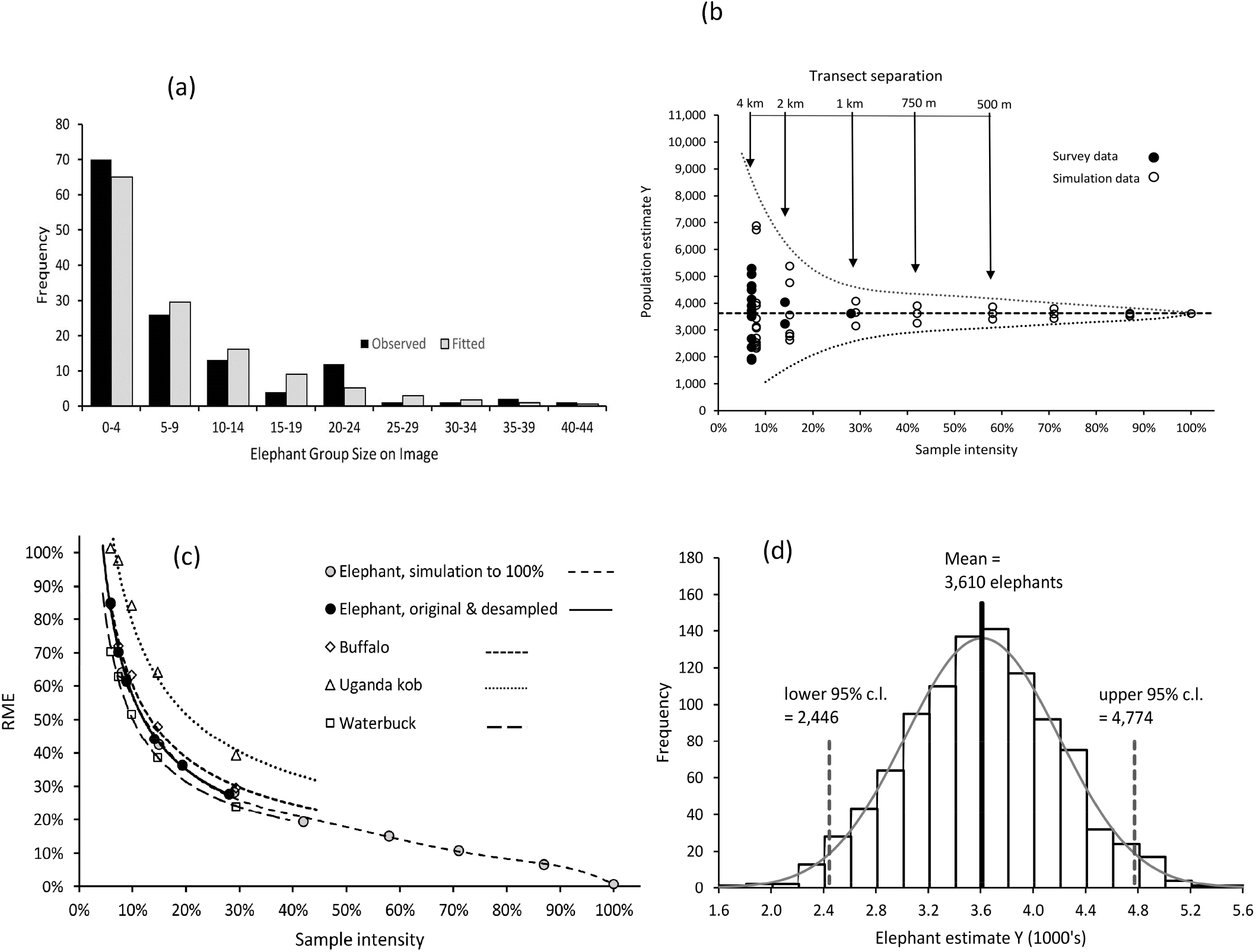
(a) For North Block, modelling the elephant group size distribution with a gamma distribution, (b) effect of transect separation (and hence sample intensity) on population estimate and highest/lowest 95% cl (dotted line), (c) RME as a function of sample intensity, (d) calculating 95% confidence intervals of the elephant population estimate using a 1000-iteration bootstrap.

Figure 8(b) shows the effect of de-sampling transects on the Jolly II estimate Ŷ for North Block, below the implemented 28% intensity, and upwards to 100% using the simulated population. The graph shows the maximum and minimum confidence limits possible from drawing any combination of regularly separated transects in a systematic reconnaissance flight (SRF) pattern, ie aircraft flying transects 1, 3, 5, 7… (14% sample) or 2,4, 6, 8… (also 14% sample). The conclusion is that, for a standard elephant counting block of 1000 sq.km, elephant sampling below an intensity of 28% is relative useless for generating an RME of ≤ 20%, ie within the recommendations of CITES-MIKE (2019). Furthermore, with a simulated total of 3 610 elephants for North Block (slightly less than the Jolly II estimate), it is also noteworthy that estimates rapidly converge more rapidly towards the ‘true’ number than the precision estimates suggest; by 28% sample, we see little variation in the actual population estimate. These findings generally accord with those of Ferreira & Van Arde (2009) who simulated ‘known’ populations and varied the sample intensity and size of survey area to determine that accuracy of estimates generally improves ahead of any increase in precision, especially when animals are clustered.

Figure 8(c) shows the improvement of precision through increasing sampling intensity up to 28% for buffalo, kob and waterbuck, and increasing to 100% sample intensity with elephant. Note that RMEs for waterbuck, which occur in small groups, and are widely dispersed, indicate more precise counts for this species than for elephants at the same sample intensity. The converse is true for kob, which occur in localized aggregations around leks; here a 28% sample returns a less-precise high RME of 40%.

Figure 8(d) shows the results of a 1000-iteration bootstrapping the Jolly II estimates of the implemented 28% survey, by randomizing the selection (by replacement) of the 38 transects across North Block. The 95% confidence limits using the percentile method are indicated as 3 610 ± 1164 (95%.cl), which is close to the Jolly II estimate of 3611 ± 1056 (95% c.l) indicating that precision can also be determined using bootstrapping. Finally, the Marriot Method returns the estimate of 3 610 ± 1210 (95% c.l.), which provides lower precision than the standard Jolly II procedure, and therefore for elephants offers no advantage.

Figure 9 indicates the statistical power of various sampling intensities for determining future elephant changes in QEPA, with effect size 40%, 60% and 80%. Figure 9(a) indicates the probability p of a Type 1 error at different sample intensities with examples indicated at *α* = 0.05 and 0.1. Figure 9(b) indicates the power of the test, with probability of a Type II error at *β* = 0.2 and experimental power *1-β* = 0.8. For example, to measure an elephant population change of 40%, either up or down, at *α* = 0.05, a 38% sample is needed. If a Type II error is to be avoided at *β* = 0.2, the sample intensity should be increased to 57%. This demonstrates that flying strip samples with 2 × 150 m width at the previously-used 2.5 km transect spacing, generating a sample intensity of 12%, is not sufficient to reliably detect even an 80% change (‘effect’) in QEPA’s elephant population. At the same time, there is no possibility to avoid a Type II error at *β* = 0.2.

**Figure 9.**
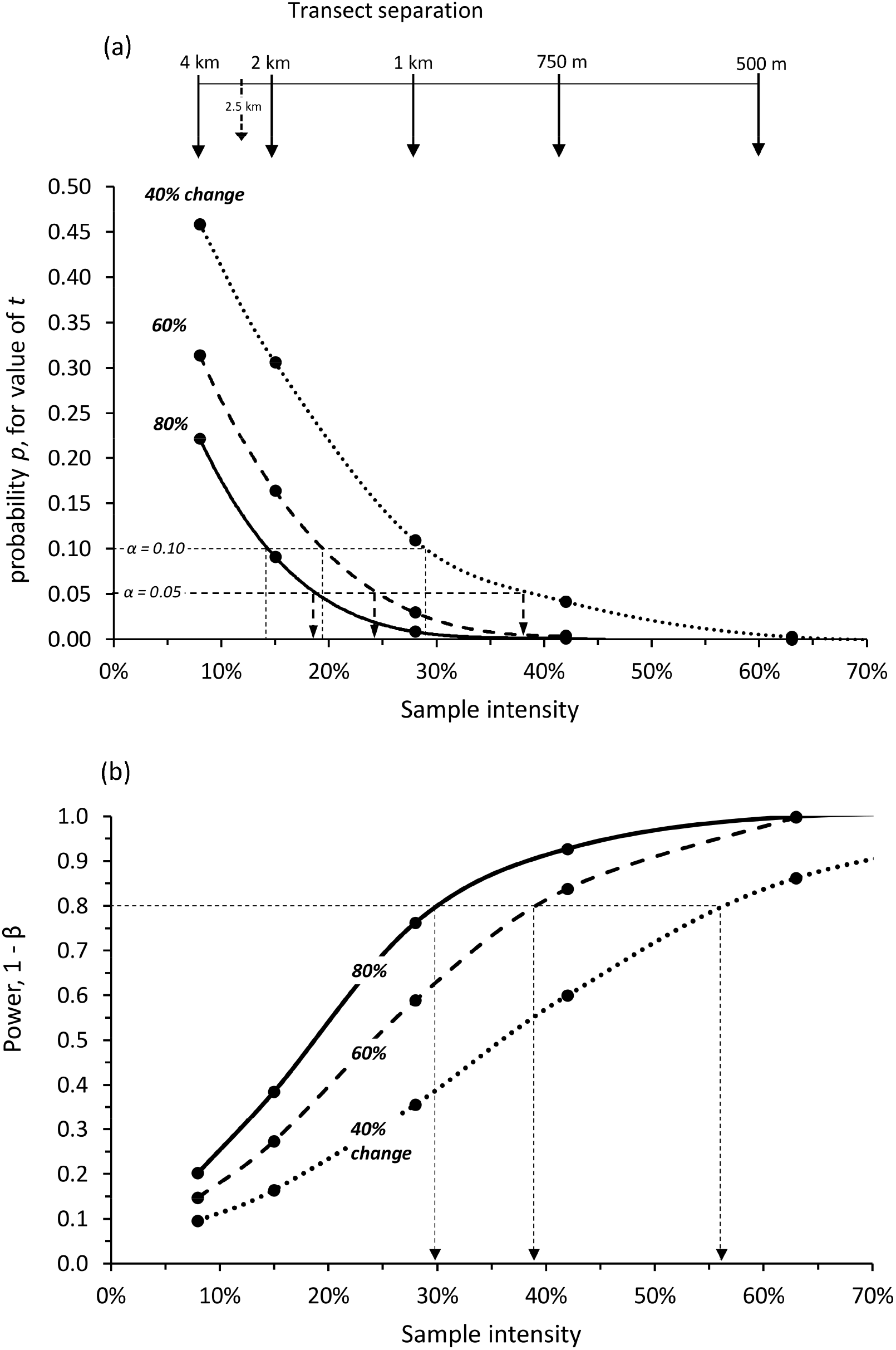
Retrospective power analysis of measuring population changes (‘effects’) of 40%, 60% and 80% of the QEPA North Block elephant population, where (a) we determine sample intensity to avoid a Type I error at *α* = 0.05 and 0.10 and (b) we determine sample intensity to avoid a Type II error at *β* = 0.2; the power of the test is given by 1-*β*. For further explanation see text.

Only by sampling at > 25% intensity, as we have done in this survey (at 28%), will generate a significant result at *a* = 0.05; this would be sufficient to detect an effect of >60% change, but insufficient for 40%. If we wanted to avoid a Type II error in measuring an effect of 60%, we would have to increase the sample intensity to 38% or 600 m transect separation, clearly an expensive proposition. The only way to improve this situation is to increase the width of the strip, which is a possible by changing the focal length of the camera lenses to capture, for example, a strip of 250 - 300 m rather than 150 m.

### 3.4 Buffaloes and hippopotamus

Buffalo are widely distributed throughout QEPA (except Dura), with the large concentrations in open grasslands and around and waterholes in Ishasha, Kyambura, Kasenyi Plains and south of Kazinga Channel, see Figure 10(a). The estimate of 14 616 ± 1974 (SE) indicates an increase in buffalo since the 1980s and 90s, when the estimates stood at around 5000 (Lamprey & Michelmore, 1995). The buffalo population is slightly lower than the estimates of 18000 counted in 1968-69 using aerial photography of large herds (Eltringham & Woodford, 1973).

**Figure 10.**
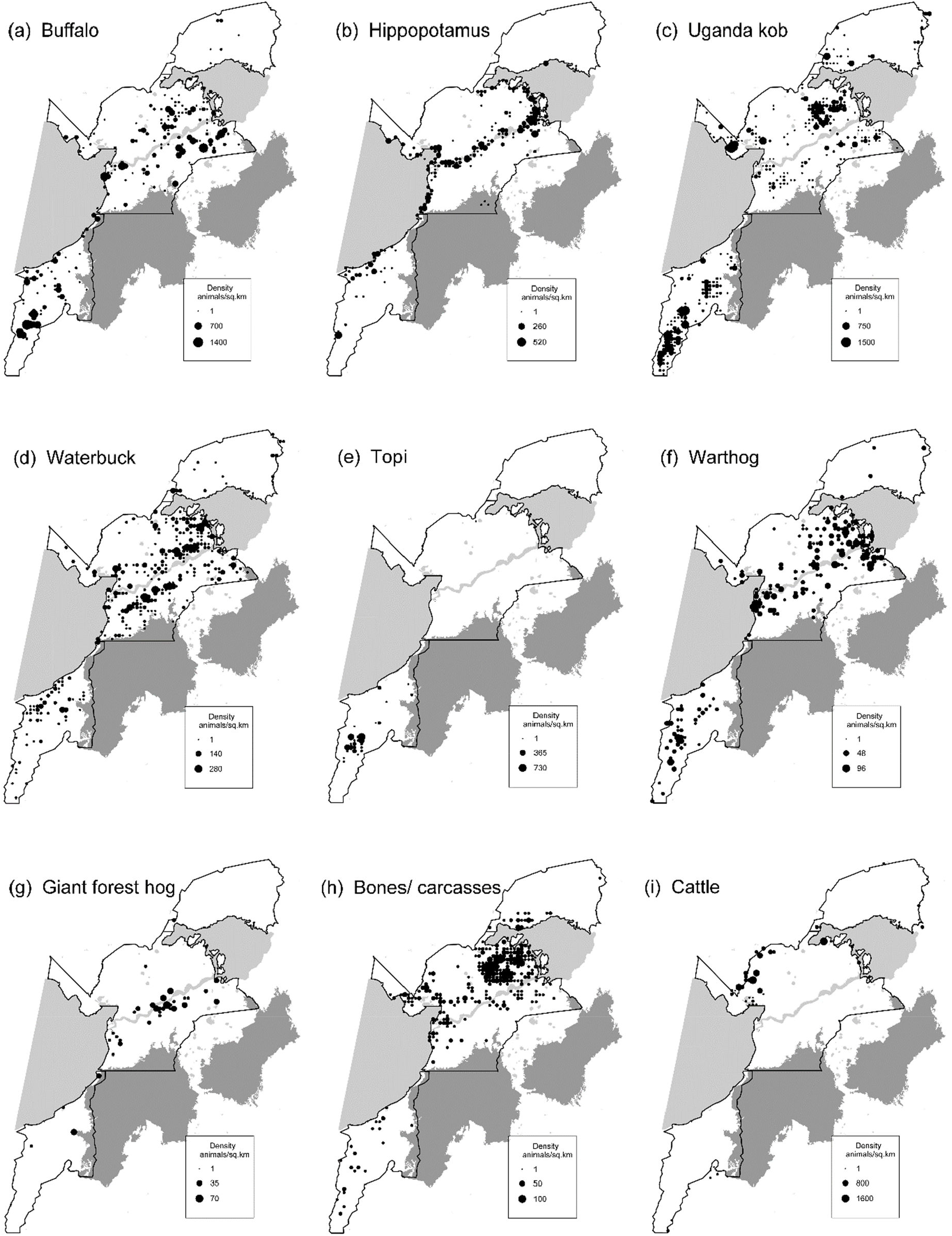
(a)-(h). Distribution and density of large wildlife species in QENP.

The hippo population is estimated at 5 716 + 585 (SE), with the main concentrations along the Kazinga channel and its confluences with Lake Edward and Lake George, see Figure 10(b). Relatively few hippos were encountered in Ishasha block; here there are anecdotal reports of intensive hippo poaching by Congolese gangs along the shoreline of Lake Edward. Overall, the 2018 estimate is higher than the 2014 GEC survey estimated 4155 + 1164 (SE), this probably being due to RSO-detection factors for the GEC count.

### 3.5 Antelopes

The estimated population of Uganda kob in QEPA is 18613 + 4496 (95% c.l). For North Block, with Jolly II population estimate of 9290 ± 3235 (95% c.l), the Marriott method greatly improving the RME from 35% to 24% (9290 ± 2201, 95% c.l) indicating that the method is useful for improving precision for heavily clumped distributions. The main kob ranges are the Kasenyi Plains in the north, Pelican Point in the west and Ishasha Plain in the south, see Figure 10(c). These distributions are similar to those recorded in all RSO counts going back to 1995. With improved law enforcement in Dura, kob are now recolonizing the area through the small corridor at Muhokya to the west of Lake George. Very few kob were encountered in the former lek areas between Craters and the Kazinga Channel, probably due to the increase in both bushland cover and *Imperata* grassland in this area.

Waterbuck are found in highest densities in the Kasenyi Plains, Kazinga South and the wetland areas east and north of Lake Edward Flats, see Figure 10(d). The survey estimate is 5 826 ± 582 (SE). This is more than double the previous aerial estimates of 2500-3500 (Wanyama et al., 2014). Waterbuck are cryptic and manyare missed in RSO-based aerial counts. In ground-counts in a range of habitats in the late 1960s, Spinage (1970) estimated an average density of 2.1 waterbuck / sq.km, or about 5000 for the park, whilst Eltringham and Din (1977) estimated a total of 4400. With high variation in waterbuck estimates over the last 50 years, no clear trend can be discerned.

The survey confirmed the ‘traditional’ localized distribution of topi in QEPA, which are found on the Ishasha Plains, with the largest concentrations close to the sector HQ, see Figure 10(e). The 2018 population is estimated at 1550 ± 583 (SE). Previous surveys indicated a decline in topi in QEPA from 5000 in the 1960s (Eltringham & Din, 1977; Jewell, 1972) to about 3000 by the mid-70s (Yoaciel & Van Orsdol, 1981), and just 100 in Year 2000 (possibly due to emigration to Virunga) (Lamprey, 2000). Since then, there has been a steady increase to the present level of 1500-2000 animals (Wanyama et al., 2014), and as confirmed by this 2018 survey.

### 3.6 Warthog and giant forest hog

The estimate for warthog is 2451 ± 261 (SE). Warthog, once numerous around Mweya and the game-viewing circuits along the north bank of Kazinga Channel, have declined significantly in these areas since 2006-2014 counts, and there are reports of intensified high predation by lions in these areas. At the same time, they have increased around village areas, especially Kazinga, Kasenyi and Katwe villages; this might be a response to this predation. The distribution is shown in Figure 10(f). No clear trend in warthog numbers is discernible, although the overall 2018 estimate is slightly higher than in the 2006-2014 counts, which have estimated 1300-1500 warthog.

The 2018 count is the first attempt to record giant forest hog from the air. The population estimate is 346 ± 83 (SE). Forest hog were once common on the Channel track area along the north bank of Kazinga Channel, but in the survey only one observation was made in this area. The highest congregation is in the thickets directly around the park HQ at Katunguru, with other isolated observations in Kyambura WR, Kisenyi, and east of Rwenshama, see Figure 10(g). Although a large proportion of giant forest hogs will have been missed in dense vegetation, the survey highlights the extremely localized distribution of this species, especially in ‘safe areas’ such as immediately adjacent to the park headquarters at Katunguru.

### 3.7 Small animal carcasses

The OCC count mapped bone sites and carcasses, these being distinguished from elephant carcasses by lack of the defining skull. As indicated in Figure 10(h), these occur at particularly high density in the Kasenyi Plains and Dura South Plains, and with lower but significant densities around Katwe town and Lion Bay. Carcass visibility will be higher in heavily grazed short-grass plains, but it is unusual that in the Ishasha Sector with its high wildlife densities, carcass densities are much lower. Clearly, there are high levels of animal mortality on Kasenyi plains, an area that is periodically heavily grazed by livestock from Hamukungu and Kisenyi.

### 3.8 Livestock

We estimated 5857 ± 1394 (SE) cattle within the protected area, or on the immediate boundaries. As shown in Figure 10(i) and recorded elsewhere, the main cattle incursion is into the Craters area from the Basongora pastoral communities living beyond the Nyamagusani River. Goats within QEPA, estimated at 2316 ± 486 are associated with individual households of the community enclaves.

### 3.9 Vegetation mapping

The vegetation classes were mapped out on a 2 km grid for Phase 1 imagery, see Figure 11. The mapping of the major vegetation types closely accords with recent (2006) mapping by satellite imagery (Jaksic-Born, 2009; Plumptre, Kirunda, et al., 2010), with the difference that the cover of thickets has greatly increased along the Kazinga Channel, with extensive spread alongthe southern side of the channel in Kyambura Wildlife Reserve. A particular strength of OCC vegetation mapping is the ability to differentiate the invasive grass species *Imperata cylindrica* (Cogon grass) from *Themeda triandra* and *Hyparrhenia* grasslands. The survey indicates that over half of the open grasslands of QEPA have now been overrun by Cogon grass. The most significant feature is the increase of *Imperata* grassland in the Craters area west to the Nyamugusani River, and south of the Kazinga Channel. Both areas have low densities of kob, waterbuck and buffalo.

**Figure 11.**
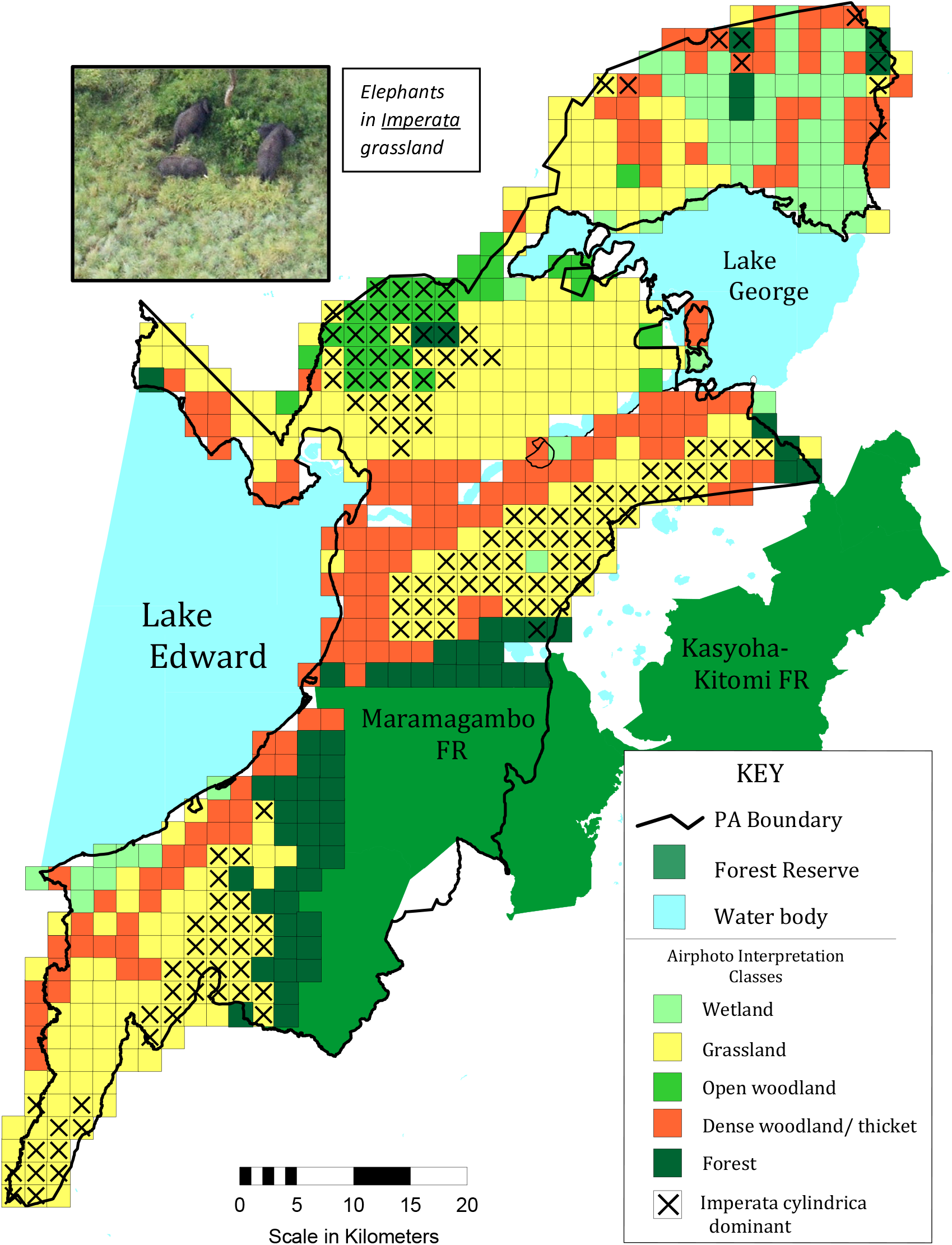
Vegetation of QEPA, as interpreted from the OCC aerial imagery. The interpretation is from the Phase 1 survey, with plotting to a 2 km grid. Areas marked as crosses indicate where *Imperata cylindrica* is the dominant grass species.

## 4. DISCUSSION

### 4.1 Trends in large wildlife species

After 40 years of ‘grey literature’ survey reports, this paper is the first to publish wildlife population trends, updated wildlife population estimates, and distribution maps in QEPA since the wildlife decimations of the late 70s and early 80s (Douglas-Hamilton et al., 1980). The 2018 survey recorded higher wildlife estimates than those obtained in previous RSO-based counts conducted since the turn of the millennium. QEPA’s biomass, which was Africa’s highest in the 1960s at approximately 19000 kg.km^−2^ (Coe et al., 1976; Petrides & Swank, 1965), and which dropped to 6500.kg.km^−2^ in the 1990s (Lamprey, 2000; Lamprey & Michelmore, 1995; Rwetsiba, Lamprey, Tumwesigye, & Aleper, 2002), now stands at approximately 14400 kg.km^−2^. This is roughly equivalent to that of the Serengeti as calculated using data of Hopcraft et al. (2015) with species weights of Coe et al. (1976).

The elephant population of 4711 ± 1106 (95% c.l), is now the highest ever recorded estimate for QEPA. The QEPA North Block elephant density of 3.4 elephants.km^−2^ is now exceeded only by Lake Manyara NP in Tanzania (Douglas-Hamilton, 1973) with 5 elephants.km^2^, and by just 5 of the approximately 190 elephant counting blocks of the vast Kavango-Zambezi Transfrontier Conservation Area of southern Africa with maximum of 8 elephants.km^2^ in Shapi Block, Hwange NP in Zimbabwe (GEC, unpubl.data).

With effective conservation measures, poaching in QEPA is at a low level as reflected in the low carcass ratio of just 1.4%. Whilst our elephant densities in peripheral areas suggest a possible ‘invisible’ Maramagambo population of 1400 elephants, other research in forest areas would put this estimate lower; Wing & Buss (1970) in the 1960s estimated that the adjacent forested Kibale NP supported 0.85 elephants.km^−2^, which would suggest some 400 elephants within Maramagambo. Through dung counts, UWA estimated some 487 elephants in Kibale in 2010, giving a similar density of 0.64 elephants.km^−2^ (Wanyama, 2010).

Our 2018 survey increases Uganda’s national elephant population estimate by 35% (UWA, 2015). This increase is likely due to three factors. The first reason is that improved aerial camera-counts have improved detection of elephants in the dense thickets of the Kazinga Channel and on the edges of the Maramagambo Forest. Elephants may have been present in similar numbers in previous RSO-based counts, but they were not detected. Clearly, camera-count methods offer great advantages for elephant counting in densely wooded areas, in that they enable a ‘freeze’ of the counting strip for focused counting. Images reveal elephants in dark forest and woodland shadows. Herds under canopy cover, invisible from one camera angle, are visible from another in frame overlaps. Partially hidden animals can be detected from glimpses of tusks or ears; and small calves can be seen beneath adults.

Secondly, protection measures by UWA are greatly intensified over the last 20 years. Ranger patrols in QEPA have extended to new areas, parks infrastructure improved, and tourism has greatly increased, although recently these efforts have been constrained by Covid-19. Recently, UWA, working in partnership with the Uganda Conservation Foundation (UCF) has constructed some 12 new ranger posts across the protected area to expand the area of patrol coverage from 10% to 70%.

Thirdly, the counting difference might reflect immigration of elephants from DRC, but in earlier PNV surveys of 2010 so few elephants were seen in PNV that this is not likely to be a major factor. Recently, a large group of elephants numbering 500+ was observed in a patrol flight in PNV, these elephants having crossed from QEPA (Alberts, 2020). The elephant groups in QEPA are relatively small, indicating general low levels of harassment and poaching, and therefore the observed very large herd in PNV might represent a ‘protective aggregation’ as QEPA’s herds expand their range back into former areas.

With severe agricultural encroachment into PNV west of Lake Edward, the movement of elephants between the central African Congo rainforest and QEPA-PNV is constrained to two extremely threatened corridors, one of just 2.5 km width at the rapidly-expanding village of Kayanja in QEPA (see Figure 1), the other in PNV of 4 km width linking the Park to the Ituri Forest in the north; the latter corridor area, near the town of Beni, is beset by rebel militia groups. These two small corridors, existentially important for biodiversity connectivity of the Greater Virunga Landscape, are amongst the vulnerable in Africa. Without their protection, QEPA-PNV will soon become ecologically isolated from the central African rainforest.

The 2018 survey also updated the status of other species in QEPA. Historically, the lakeshores and associated vegetation of QEPA have been altered by heavy grazing and trampling by the world’s largest hippo population. In the late 1950s hippo numbers in QENP were recorded as 14000+ (Bere, 1959), and a culling programme was instigated to reduce severe overgrazing on Mweya peninsular and along Kazinga Channel. Over the 1960s, some 1000 hippos were culled per annum, with the result that the vegetation recovered in these areas, and diversity of other species increased (Field, 1970). It is not clear how many hippos remained by the late 1960s, but it is suggested that they will still relatively numerous, if wary (Field & Laws, 1970). Hippos were counted by aerial sample count in 1980 (Douglas-Hamilton et al., 1980) and 1995 (Lamprey & Michelmore, 1995), but since then UWA has counted hippos by boat, with participation of tourism groups.

The main areas for hippopotamus are Lion Bay, Kazinga Channel and the confluence of the latter with Lake George. Hippo numbers in the Ishasha sector of QEPA are significantly reduced, anecdotal reports indicating that there has been intensive poaching by hunting gangs operating from the adjacent DRC. Hippo meat is much favoured by local communities around QEPA, with studies in 2007-8 indicating some 280 hippos killed per year (Olupot, McNeilage, & Plumptre, 2008), contributing some 90% of bushmeat biomass to communities. The 2016 UWA boat-based hippo counts in the Lakes and Kazinga Channel indicate a population of 6500 hippos; our 2018 aerial estimate of 5 716 hippos reflects high visibility in the OCC count, and the ability to count hippos not just in the lakes but in the waterholes in dense thickets. Applying a correction factor of 1.3 for hippos underwater at the time of aircraft overpass (Mackie, Dunham, & Ghiurghi, 2013) would bring this total to about 7 400 hippos for QEPA.

The Uganda kob, buffalo and topi populations are now estimated at 18613, 14616 and 1550 respectively. The overall trend suggests that in the earlier 2014 RSO-based count (Wanyama et al., 2014) they were well detected and enumerated by observers. Therefore, although localized clumping of animals within parts of the survey area leads to low precision, the aggregation of animals into large herds in open areas that are easily detected by RSOs and photographed can result in improved detection (Schlossberg et al., 2016). Paradoxically, for the 2018 QEPA count the possibility also exists that enhanced camera detection actually shows a reduction of kob, since RSO-based counts are shown to miss as many as 50% of this species (Lamprey et al., 2020). Meanwhile, for cryptic animals that in occur mainly in small groups of 1-5 animals, such as waterbuck and warthog, RSOs in previous counts may have missed many.

No clear statement can be made about the low RSO-based buffalo and kob estimates of 2010 that caused concern in QEPA’s management; it is possible that these species were subjected to a severe stint of poaching from which they recovered quite rapidly by 2014 − a conclusion reflected in anecdotal management reports. In our 2018 camera-count, these populations are at levels that are significant, but are well below the high point of the 1990s. In Dura Block, an area close to the urban centre of Kasese with high bushmeat demands, recent wildlife protection measures have resulted in an increase in Uganda kob and buffalo numbers after years of intensive illegal hunting in the area. However, in their main areas on the Kasenyi Plains, kob and buffalo populations are stagnating. The high density of carcasses on the Kasenyi plains points to high wildlife mortality, possibly linked to compression of range by the spread of *Imperata cylindrica*, continued anthrax in the populations and poaching.

This study, and the trial that preceded it in 2015 (Lamprey et al., 2020), is the first known example of using a systematic aerial survey for determining carcass ratios for animals other than elephants. Whilst bones of smaller animals will probably be visible for a year before disintegrating fully, the inclusion of this category enable comparisons to be made with other areas. In a test stratum of the Kasenyi Plains, the area with the highest kob density in QEPA, the carcass ratio (Douglas-Hamilton & Burrill, 1991) is calculated at 14% of the kob and buffalo population, with a density of 16 bone patches, km^−2^. For comparison, in a similar open area of Buligi in Murchison Falls NP, where kob have been increasing at 12% per annum since Yr 2000, the carcass ratio for large mammals (also including hartebeest and giraffe in this environment) is just 1%, with 1.6 bone patches.km^−2^. Predation can be ruled out as a factor of this high mortality, since the number of lions in both areas remains low at about 140 lions in QEPA and 130 in Murchison Falls (Omoya, Mudumba, Buckland, Mulondo, & Plumptre, 2014).

This count was innovative in determining the extent of an invasive plant species. *Imperata cyllndrica*, long-recorded in QENP (Langdale-Brown et al., 1964; Lock, 1993), has a distinctive tussock growth form easily identified on low-level oblique aerial photographs (see Figure 11). Our survey revealed that it is spreading very rapidly and now occupies some 50% of QEPAs grassland areas. Indigenous to Asia and parts of Africa, *Imperata* is invasive and regarded as a serious weed due to root system effects on germination of other species, and difficulties in eradication (Holm, Plucknett, Pancho, & Herberger, 1977). *Imperata* is fire-tolerant, and the spread appears to be the result of fires when herbivory has been reduced, as was experienced in the 1970s. Once established, increased fire frequency results in further spread (Plumptre, Kirunda, et al., 2010). When it has reached tussock form, it becomes unpalatable and its spread in QEPA has now excluded Uganda kob from previous grazing and leking areas (Jaksic-Born, 2009).

### 4.2 Improving accuracy and precision using cameras and intensified sampling

Following earlier tests of aerial camera methods in Murchison Falls in Uganda, and Tsavo and Ol Pejeta in Kenya, the QENP aerial camera count represents the next stage of operationalization of oblique-camera-counts for estimating wildlife populations. The count broke new ground in QEPA in other aspects of aerial sampling. Firstly, in comparison with previous RSO-based counts, it was successful in generating a two-fold improvement of precision in population estimates; over the entire protected area, we achieved a relative margin of error (RME) of 23% for elephants, and 20-27% for the other major species. This was obtained with one of the highest elephant densities in Africa at >3 elephants.km^−2^, sampled at 28% intensity whilst adhering to sample strip parameters indicated in international counting standards (CITES-MIKE, 2019).

In the context of sample intensity, it is useful to compare our North Block (1120 sq.km) sample strategy with similar survey areas in Africa. In the Susuwe Block of Namibia (1123 sq.km) with an elephant density of 2.4 elephants/ sq.km, a 30% sample achieved an RME of 46.7% (Gibson & Craig, 2015). In Chipinda Pools (1075 sq.km) in Gonarezhou, Zimbabwe, with 2.7 elephants sq.km sampled at 22%, Dunham et al (2013) obtained an RME of 34%. In Botswana’s Okavango, for a survey area of 1200 sq.km (Vumbra, Duba, NG11 blocks) sampled at 20%, Schlossberg et al (2016) derived an RME of 35% for an elephant density of 3.2 elephants/sq.km. These results all suggestthat the target RME of < 20% specified by CITES-MIKE (2019), is essentially unachievable under the guidelines for relatively small areas such as QEPA. Ferreira & Van Arde (2009) showed that RME is a function not only of population density and sample size, but of area size, and thus the aggregation of sample blocks can improve precision.

This survey demonstrates imaging technologies that can be used to enhance precision by increasing the sample size at no extra cost. Currently, the 150 m sample strip as recommended by CITES-MIKE (2019) derives from experiments in the 1960s and 1970s where it was found that, at standard SRF flying height of 300 ft above ground, the observers ability to detect animals decreased with increasing strip width (Caughley, 1974; Norton-Griffiths, 1978; Pennycuick & Western, 1969); by the 1980s the 150 m strip had become a standard in SRF methodology (Craig, 2012; Ottichilo & Khaemba, 2001; Stelfox & Peden, 1981). This limitation constrains sample intensity, as demonstrated in our survey. Today DSLR and mirrorless cameras are available with over 60 MP pixel density. These can befitted with wider-angle lenses, or the aircraft platforms can be flown higher, enabling an increased strip width and higher sample intensity. This increased sample intensity can be achieved whilst retaining the necessary ‘ground-sampling-distance’ (GSD) of < 5 cm for detection of smaller animals. For example, in QEPA, we could have increased our sample intensity to 60% merely by flying the aircraft at 1200 ft rather than 600 ft; keeping the same 24 MP cameras would have increased the GSD from approximately 2.5 cm to 5 cm.

A third improvement relates to the flight parameters. The QEPA count was conducted at a slightly higher altitude (600’ HAGL) than the previous 500’ HAGL used in Uganda (Lamprey et al., 2020) to reduce, at least notionally, the possibility of the aircraft being hit by a stray AK47 bullet; at the time of the survey there was armed cross-border conflict overfishing rights in Lake Edward. The higher altitude, offset by longer focal length of the camera lenses, proved equally effective for camera-count surveys, and has been adopted in more recent OCC surveys. This can increase the safety of the survey mission, especially when operating in rugged and hilly terrain.

### 4.3 Potential for machine learning

The interpretation of some 43000 images in QEPA required the efforts of 5 airphoto interpreters working fulltime over 2 months. Most of these images were True Negatives (TNs), that is images with no target species, and therefore much labour was expended on eliminating these images. In aerial imagery gathered by systematic sampling, the proportion of True Positive (TPs) animal images − those with confirmed animal(s) − in the entire image dataset depends on the environment. In the highly productive and well-watered protected areas of Uganda, with their abundant wildlife populations, some 10% of images have at least one target animal. In the arid environments of Kenya, this proportion is just 2% (Lamprey et al., 2019).

New strides in machine learning with convolutional neural networks such as RetinaNet and Faster-R-CNN have the potential to accelerate the counting of animals on aerial imagery through elimination of True Negatives and through species-level animal detection (Duporge, Isupova, Reece, Macdonald, & Wang, 2021; Kellenberger, Marcos, & Tuia, 2018). Training of the CNNs requires massive image annotation by humans (Eikelboom et al., 2019). Recently, there has been particular success in identification of large and/ or conspicuous species, such as elephant, giraffe, zebra in Kenya (Eikelboom et al., 2019) and kiangs (Asiatic wild ass) of the Tibetan Plateau (Peng et al., 2020). Closer to our study area, Delplanque et al (2021) developed CNN algorithms to correctly identify 73% of elephants, buffalo, Uganda kob and topi on UAV imagery in Virunga and Garamba National Parks in DRC. In all of these machine learning studies, the primary objective is to eliminate as many True Negatives as possible, whilst also keeping the False Positives (FPs) − incorrectly flagged targets − to acceptable limits. Currently, the best FP/TP ratio in multi-species stands at about 1.5, meaning that for every 2 correctly identified animals, there are 3 incorrectly identified ones (Delplanque et al., 2021; Eikelboom et al., 2019). However, if at this stage ML is considered as a tool to reduce human interpreter workloads, any solution that reduces 40% or more of the work without compromising accuracy is good news (Kellenberger, Marcos, Lobry, & Tuia, 2019). Much more insidious are the False Negatives, because if the population of a species is low, the failure of a CNN to detect even a few individuals has a large impact on population estimates.

Currently the graphics processing unit (GPU) costs of using web-based CNN platforms such as Azure or Google Labs to the filtering of billions of pixels acquired in an aerial camera count of eg 40000 images exceeds, by a factor of at least three times, the cost of employing Ugandan postgraduate students in interpreting the imagery. These technical challenges will, in time, be addressed through ongoing developments and the costs of processing pixels are expected to fall.

## Data Availability Statement

The datasets used in this analysis are gathered in collaboration with Uganda government agencies. Due to their sensitive nature in wildlife management, they are not made publicly available. However, requests for raw data for research purposes will be considered on a case-by-case basis.

## Acknowledgements

We appreciate the comments of Dr lain Douglas-Hamilton and Dr Chris Thouless of Save the Elephants in an early draft of this manuscript. Jerry Burley, Patrick Agaba and Derek Lubangakene of UCF provided great logistical assistance in the field. Our thanks to Bernadette Apio, Dorcus Ninsiima, Innocent Kasekendi, Conslate Alezuyo and Juliet Kyakobyewo for their hard work in analysing the thousands of images.

